# Multiscale Characterization of Complex Binding Interactions of Cellulolytic Enzymes Highlights Limitations of Classical Approaches

**DOI:** 10.1101/2020.05.08.084152

**Authors:** Shishir P. S. Chundawat, Bhargava Nemmaru, Markus Hackl, Sonia K. Brady, Mark A. Hilton, Madeline M Johnson, Sungrok Chang, Matthew J. Lang, Hyun Huh, Sang-Hyuk Lee, John M. Yarbrough, Cesar A. López, S. Gnanakaran

## Abstract

Cellulolytic microorganisms, like *Trichoderma reesei* or *Clostridium thermocellum*, frequently have non-catalytic carbohydrate-binding modules (CBMs) associated with secreted or cell surface bound multidomain carbohydrate-active enzymes (CAZymes) like cellulases. Mostly type-A family CBMs are known to promote cellulose deconstruction by increasing the substrate-bound concentration of cognate cellulase catalytic domains. However, due to the interfacial nature of cellulose hydrolysis and the structural heterogeneity of cellulose, it has been challenging to fully understand the role of CBMs on cellulase activity using classical protein-ligand binding assays. Here, we report a single-molecule CAZyme assay for an industrially relevant processive cellulase Cel7A (from *T. reesei*) to reveal how subtle CBM1 binding differences can drastically impact cellulase motility/velocity and commitment to initial processive motion for deconstruction of two well-studied crystalline cellulose allomorphs (namely cellulose I and III). We take a multifaceted approach to characterize the complex binding interactions of all major type-A family representative CBMs including CBM1, using an optical-tweezers based single-molecule CBM-cellulose bond ‘rupture’ assay to complement several classical bulk ensemble protein-ligand binding characterization methods. While our work provides a basis for the ‘cautious’ use of Langmuir-type adsorption models to characterize classical protein-ligand binding assay data, we highlight the critical limitations of using such overly simplistic models to gain a truly molecular-level understanding of interfacial protein binding interactions at heterogeneous solid-liquid interfaces. Finally, molecular dynamics simulations provided a theoretical basis for the complex binding behavior seen for CBM1 towards two distinct cellulose allomorphs reconciling experimental findings from multiscale analytical methods.

**Significance Statement:** Multimodal biomolecular binding interactions involving carbohydrate polymers (e.g., cellulose, starch, chitin, glycosaminoglycans) are fundamental molecular processes relevant to the recognition, biosynthesis, and degradation of all major terrestrial and aquatic biomass. Protein-carbohydrate binding interactions are also critical to industrial biotechnology operations such as enzymatically-catalyzed bioconversion of starch and lignocellulose into biochemicals like ethanol. However, despite the ubiquitous importance of such interfacial processes, we have a poor molecular-level understanding of protein-polysaccharide binding interactions. Here, we provide a comprehensive experimental and theoretical analysis of bulk ensemble versus single-molecule binding interactions of enzyme motors and associated non-catalytic binding domains with cellulosic polysaccharides to highlight the critical limitations of applying classical biochemical assay techniques alone to understanding protein adsorption or biological activity at solid-liquid interfaces.

## Introduction

Plant biomass, composed of polysaccharides like cellulose, is an ideal feedstock for bioconversion into various bioproducts like ethanol (1, 2). Cellulose is a β-(1→4)-glucose polymer that self-assembles to form crystalline nano/micro-fibrils that are recalcitrant to depolymerization due to strong intramolecular hydrogen bonding and dispersion forces (3). Cellulolytic microbes (like *T. reesei* and *C. thermocellum*) have therefore evolved complex biocatalytic machineries called CAZymes like cellulases to deconstruct cellulose into fermentable sugars (4–6). Cellulases are comprised of two or more polypeptide domains called catalytic domains (CDs) and CBMs (4). CBMs tethered to cellulase CDs are characterized by a planar binding motif, that is complementary to crystalline cellulose fibril structure to facilitate cellulase activity towards insoluble and heterogenous cellulosic substrates (7, 8). While cellulases are incredibly effective biocatalysts (9), these enzymes are still inefficient for industrial applications often due to CBM-driven non-productive interactions with insoluble substrates that necessitates high protein loading requirements (4, 10).

Thermochemical pretreatment of biomass using acids, bases, or ionic liquids is therefore employed to increase polysaccharide accessibility to enzymes and reduce non-productive cellulase binding (11–13). Pretreatment with anhydrous liquid ammonia results in conversion of native cellulose I to cellulose III allomorph (14), thereby improving hydrolytic activity of several fungal (15) and bacterial cellulase mixtures (16). However, processive exocellulases such as *Tr*Cel7A (or Cel7A from *Trichoderma reesei*) and *Tf*Cel6B (or Cel6B from *Thermobifida fusca*), that are often workhorse cellulolytic enzymes, show reduced activity on pretreated cellulose III for reasons that are poorly understood (16, 17). Although the processive mechanism of Cel7A on native cellulose I has been studied extensively using classical biochemical assays (18–21) and molecular simulations (22, 23), there is still limited consensus on how to monitor initial enzyme association with cellulose chain (24) or dissociation of non-productively bound enzymes (18, 25) to identify rate-limiting factors impacting cellulose hydrolysis. Hence, there is a need to develop better experimental methods that can track cellulase binding commitment and motility in real-time with atomic-scale resolution for both native crystalline cellulose and industrially relevant pretreated cellulosic substrates.

Single-molecule fluorescence imaging has been used to determine kinetic parameters (e.g., adsorption and desorption rates) of processive exocellulases (10, 26, 27), whereas high speed atomic force microscopy imaging has been employed to track motility of single cellulase molecules (28, 29). However, these methods are unable to resolve slower sub-nanometer translational rates of processive cellulases. To address this issue, we recently reported an optical tweezers force spectroscopy-based assay to track the single-molecule motility of processive cellulase Cel7A with sub-nanometer and millisecond resolution (30). Interestingly, Cel7A CD in the absence of native CBM1 showed lower dwell times between catalytic turnover steps suggesting that CBMs could impede full-length cellulase motility possibly due to non-productive binding. This assay now provides us the opportunity to systematically track the role of CBM binding on commitment to cellulase motility. A detailed understanding of the mechanistic role of CBMs in full-length processive cellulase binding and motility on industrially relevant cellulosic substrates can enable rational enzyme engineering efforts to reduce CBM-mediated non-productive binding.

Here, we have applied our optical tweezer assay to specifically investigate the Cel7A initial binding stability and commitment to processive motility on native and ammonia-pretreated crystalline cellulose allomorphs. To further understand the role of CBMs towards cellulase binding and the observed single-molecule binding instability of Cel7A to cellulose III, we characterized the binding of two well-studied Type-A CBMs CBM1 (*T. reesei*) and CBM3a (*C. thermocellum*) using various classical protein-ligand adsorption assays (e.g., ‘pull-down’ or solid-state depletion assays). We also characterized the binding of model CBMs from all representative Type-A families to cellulose I and cellulose III using pull-down assays to generalize our findings. Since most classical pull-down assay generated data only provided ambiguous insights into the multimodal heterogeneity of CBM-cellulose binding interactions, we developed a new optical tweezers based CBM-cellulose bond ‘rupture’ assay to characterize binding of CBM1 (*T. reesei*) to several cellulosic substrates. We contrasted our pull-down assay results with findings from our CBM-cellulose rupture assay to highlight the limitations of classical approaches to gain a molecular-level understanding of binding interactions of CBMs with cellulose. Lastly, we used molecular dynamics simulations to provide a theoretical basis for the complex binding behavior seen for CBMs towards cellulose III that further supports our interpretation of the single-molecule CBM-cellulose bond rupture and Cel7A motility assay findings. In summary, our work highlights how subtle changes in CBM binding to distinct cellulose allomorphs can critically impact processive cellulase motility. Furthermore, a multifaceted approach is often necessary for characterizing the binding heterogeneity and multimodal nature of CAZyme-polysaccharide interactions.

## Results

### Single-molecule Cel7A binding and cellulase motility is impacted by cellulose ultrastructure

*Cladophora* sp. (*Cladophora glomerata*) derived highly crystalline cellulose I fibers were isolated, as described previously (30), followed by anhydrous liquid ammonia pretreatment to prepare cellulose III (31). Details about cellulose isolation, ammonia pretreatment, and cellulosic spectroscopic characterization are provided in the SI appendix methods/results section and **Fig. S1**. Motility assays were performed on both cellulose allomorphs to study how subtle structural differences impact the binding and motility of Cel7A on the single-molecule level. Details regarding Cel7A motility assay are published elsewhere (30). Briefly, Cel7A was attached via sulfo-SMCC cross-linking to the thiol tag on the end of a biotinylated 1010 bp DNA tether and attached to a 1.25 μm streptavidin-coated polystyrene bead (see **Figure 1A**). The Cel7A functionalized bead was positioned directly above a cellulose fiber to initiate binding and the bead position was monitored as the enzyme first bound, hydrolyzed, and processed along the cellulose surface for cellulose I or cellulose III fibril surface (see **Figure 1B** for cross-sectional views of model cellulose structures). Representative individual Cel7A traces and average motility for cellulose I and cellulose III are shown in **Figure 1C**. The average Cel7A velocity on cellulose I was 0.25 ± 0.35 nm s-1 (s.d.; N=68 motility traces), which is greater than that seen on cellulose III, 0.17 ± 0.14 nm s-1 (s.d.; N=30 motility traces). Reduced Cel7A motility results are consistent with our previous bulk Cel7A activity assay results reported for plant-derived cellulose III (15). However, enzymatic saccharification conducted using a *T. reesei* derived cellulase cocktail gave a >3-fold higher overall rate of enzymatic hydrolysis for pretreated cellulose III (SI Appendix **Fig. S2**). Similar improvements in cellulose III saccharification rates have been reported mostly due to enhanced endo-exo cellulase synergy on cellulose III (15–17, 31). However, as reported previously (15, 17), reduced Cel7A motility on cellulose is likely correlated with lowered bulk activity seen for processive cellulases. We hypothesized that processive cellulases like Cel7A show reduced binding and activity towards cellulose III likely due to impaired Cel7A binding and processive motility cycle initiation.

**Figure 1.**
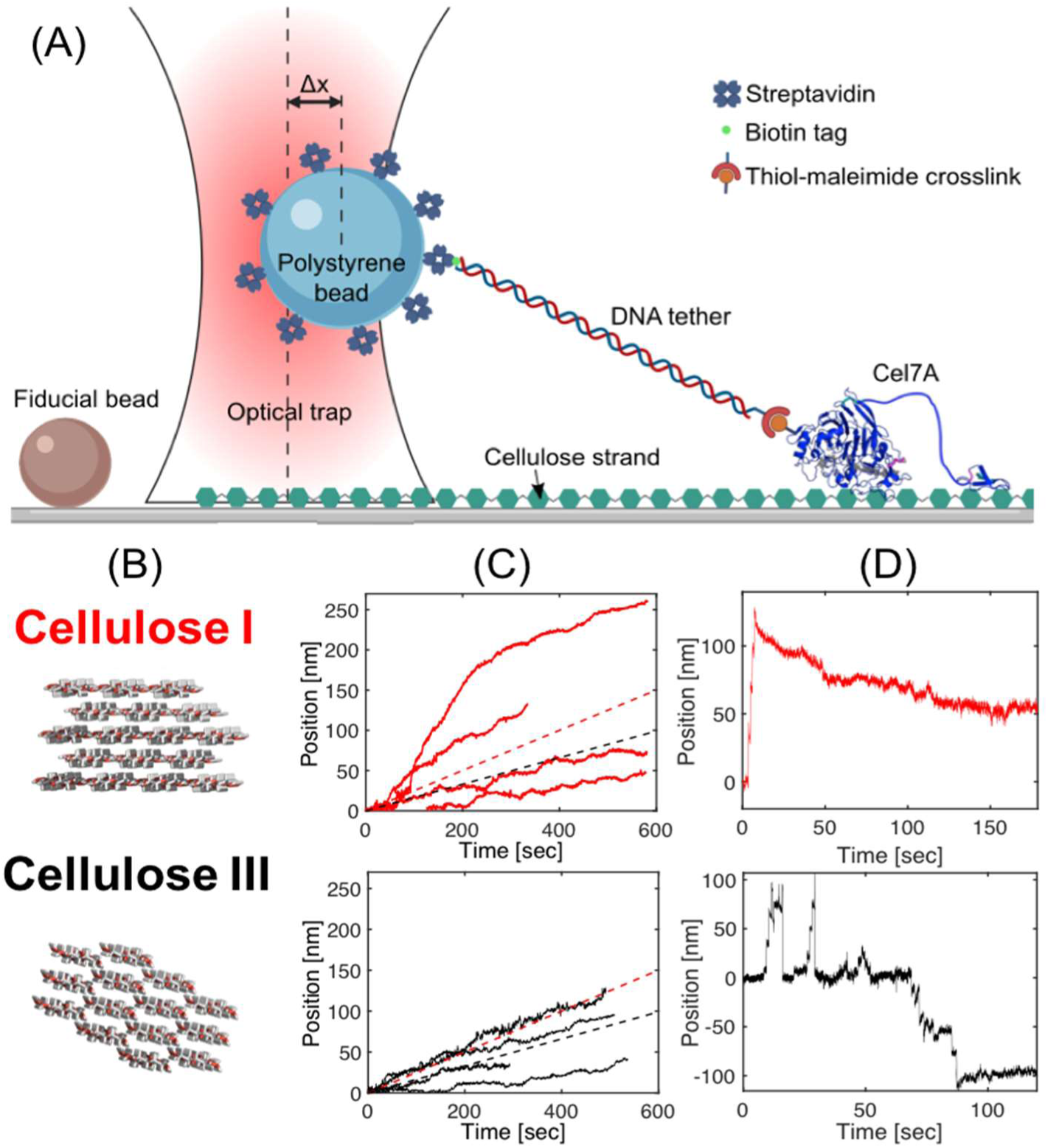
Processive cellulase Cel7A shows unstable binding and lower single-molecule motility on cellulose III. (A) Schematic of rupture assay setup (not to scale). A streptavidin coated bead is tethered to a single TrCel7A via a thiol-maleimide crosslink to a DNA linker containing a biotin tag on the opposite end. Δx represents the distance bead is displaced from trap center. Diagram created with BioRender.com. (B) Cross-sectional view of model cellulose I and III crystalline allomorphs depicting key morphological differences in fibril that arise due to distinct underlying unit cell crystal structures. (C) Representative traces of Cel7A enzyme motility on Cladophora derived cellulose I (top panel, traces in red) and cellulose III (bottom panel, traces in black) substrates are shown here. Dashed lines indicate average velocities, 0.25 ± 0.35 nm s^-1^ (s.d.; cellulose I; N=68; in red) and 0.17 ± 0.14 nm s^-1^ (s.d.; cellulose III; N=30; in black). (D) Traces representing initial stable binding to cellulose I followed by Cel7A motility (top panel; in red) and initial unstable binding to cellulose III followed by eventual Cel7A motility (bottom panel; in black) are shown here. Additional representative traces showcasing unstable protein binding prior to Cel7A motility initiation can be found in SI appendix **Fig. S3**.

During our motility assays it was also possible for us to gauge the relative commitment of processive Cel7A motion for distinct cellulose allomorphs immediately prior to motility initiation. To initiate the single-molecule Cel7A motility, a functionalized bead is positioned directly above a surface-affixed cellulose fiber and periodically pulled via the piezo stage to test for bound enzymes. Such initial binding is considered stable or committed when the Cel7A-cellulose bond survives, and the enzyme exhibits motility for a period greater than 10 s. In some cases, the full-length Cel7A was seen to bind but not commit to significant motility on the cellulose surface highlighting non-productively engaged cellulases. Alternatively, Cel7A-cellulose bond instability is revealed through initial bead displacement followed by rapid detachment. Given this criteria and observation times of 600s for each trace, Cel7A-cellulose initial engagement bond instability was determined to be lower for cellulose I (12% of traces, N=17) than cellulose III (23% of traces N=13). **Figure 1D** and **Fig. S3** show representative traces of binding stability/instability for Cel7A binding to cellulose I and cellulose III. We hypothesized that impaired Cel7A binding arises due to weaker CBM1 domain engagement with cellulose III surface and was therefore the major focus of this study.

### Type-A CBMs display higher binding affinity towards native crystalline cellulose I allomorph

Firstly, solid-state depletion or classical ‘pull-down’ binding assays were used to characterize equilibrium binding interactions for two well-studied Type-A CBMs; CBM1 (*T. reesei*) and CBM3a (*C. thermocellum*) (8). To generalize our findings further, we characterized binding interactions of representative CBMs from all major Type-A CBM families (32). Type-A CBMs bind crystalline cellulose through mostly hydrophobic stacking interactions between conserved planar aromatic residues and glucosyl units of cellulose chain (32), as illustrated in **Figure 2A**. Details regarding gene sequences, cloning, expression and protein purification strategies for all Type-A CBM families tested here can be found in the SI Appendix Materials and Methods section (33). CBMs were tagged with green fluorescent protein (GFP) since fluorescence-based methods for protein detection are several orders of magnitude more sensitive than UV-absorbance spectrophotometry (34). In addition, this fusion protein construct also helps easily visualize CBM localization on cellulose fiber surfaces (**Figure 2B**) and estimate binding rate constants using alternative fluorescence based optical imaging methods (35, 36), as discussed later.

**Figure 2.**
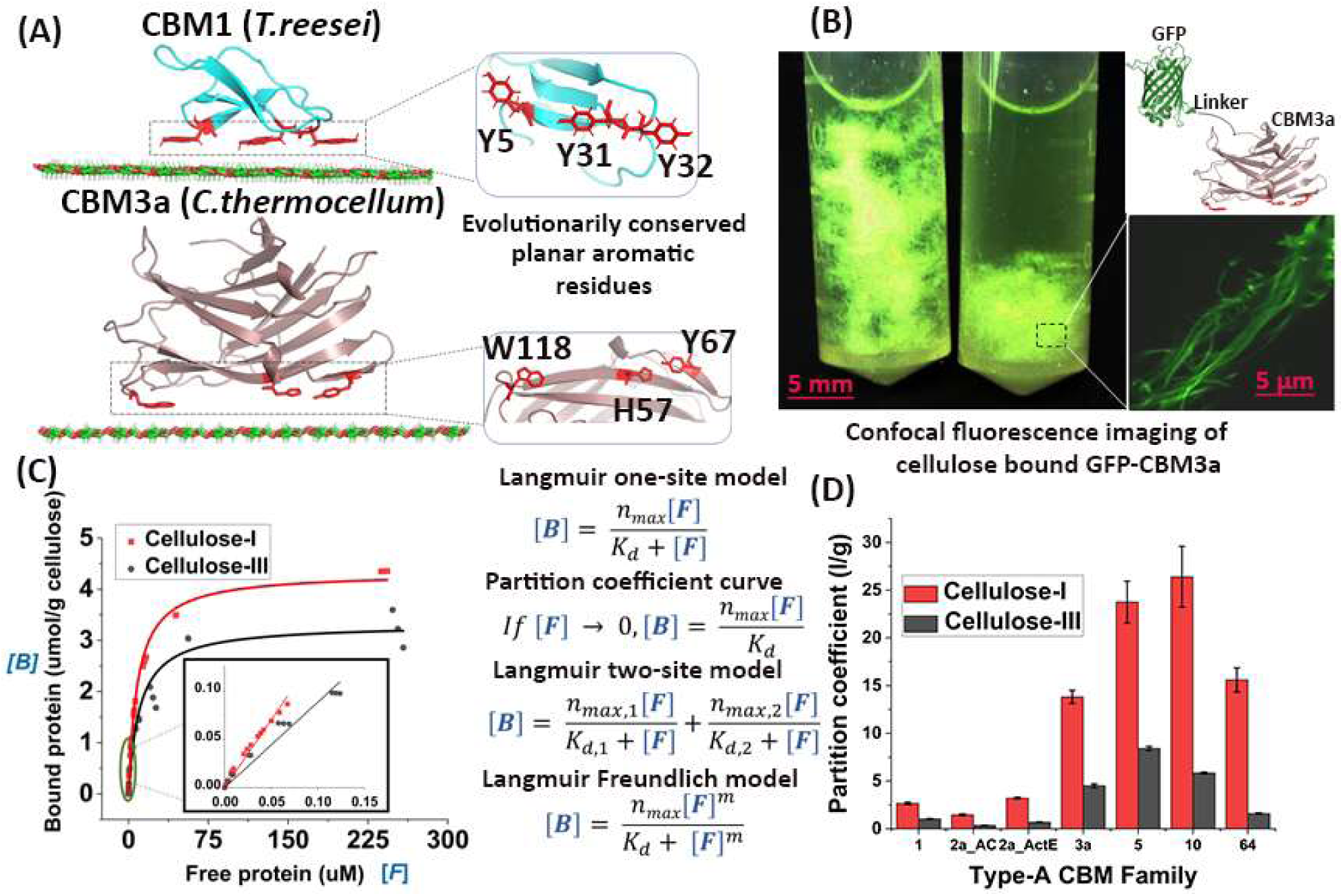
Type-A CBMs show reduced bulk binding toward cellulose III. (A) Type-A CBM1 (T. reesei) and CBM3a (C. thermocellum) docked on the hydrophobic face of crystalline cellulose. (B) Cladophora cellulose I suspension bound to GFP-linker-CBM fusion protein before (left) and after (right) ‘pull-down’ separation of cellulose bound protein fraction. Inset displays individual cellulose microfibrils bound to GFP-CBM3a. (C) GFP-linker-CBM1 (T. reesei Cel7A) equilibrium binding data for Cladophora based cellulose I and cellulose III to estimate equilibrium adsorption constants are shown here. Non-linear relationship between bound and free GFP-CBM1 concentration for cellulose I (in red dots) and cellulose III (in black dots) is shown here for replicate assays. Fitted line depicts a Langmuir one-site model. Inset graph shows the linear region of this model to estimate partition coefficient. Relationship between bound and free protein for various adsorption models such as Langmuir one-site, two-site, and Freundlich models are shown here. (D) Comparison of partition coefficients (L/g) estimated for Type-A CBMs towards Cladophora based cellulose I (in red) and cellulose III (in black) are shown here. Error bars are replicate standard deviations for reported mean values.

Full-scale equilibrium binding analyses from low to saturating protein loadings were conducted for CBM1 (*T. reesei*) and CBM3a (*C. thermocellum*), to estimate binding parameters for both cellulose allomorphs (**Figure 2C** and SI appendix **Fig. S4-S5**). Langmuir one-site/two-site and Langmuir-Freundlich based adsorption models were fitted to the protein adsorption data using non-linear regression, as described previously (7, 8, 15). The model-fitting analysis allowed estimation of the maximum available binding sites (*n*_*max*_) and equilibrium dissociation constant (*K*_*d*_), in addition to other parameters specific to the model, for CBM1 and CBM3a towards both cellulose allomorphs (**Table 1**). Depending on the fitted model, both CBM1 and CBM3a often displayed either lower or nearly comparable estimated bulk binding affinity (or larger *K*_*d*_) towards cellulose III. The *K*_*d*_ value predicted for both substrates for a given CBM family were of the same order of magnitude except in the case of high affinity sites for CBM3a when the data was fitted using a Langmuir two-site model, in which case the affinity differed by ∼13-fold for cellulose I versus cellulose III. The total number of binding sites for CBM1 to cellulose I was higher (∼1.2-1.5 fold) compared to cellulose III in all cases, except in the case of high-affinity binding sites (*n*_*max*_) for the two-site model. In the case of CBM3a, the total number of binding sites predicted for cellulose I was slightly lower than cellulose III except when the data was fitted using a one-site model. However, a closer inspection of the data (SI Appendix **Fig. S5**) suggests that even at the highest CBM3a concentrations tested (∼50 μM), proper saturation behavior might not be fully observed. These results emphasize the need for cautious use of Langmuir adsorption models fitted to CBM-cellulose pull-down assay data since the lack of high-quality data at higher protein concentrations can lead to spurious binding parameter estimation. A detailed discussion regarding the molecular basis for interpretation of these results using multi-site binding models and the shape of the Cladophora cellulose fibrils based on AFM imaging (see SI Appendix **Fig. S6**) can be found in the SI Appendix Results and Discussion section.

**Table 1.** Langmuir-based model parameters for GFP-CBM1 and GFP-CBM3a adsorption to Cladophora-based cellulose I and III. Here, binding dissociation constant (K_d_; μM), maximum available binding sites (n_max_; μmol/g cellulose), and Freundlich power constant (m) fitted parameters are shown. Standard root-mean square errors (RMSE) for the estimated best-fit parameter values are shown as well. Model fitting details for all Langmuir-based adsorption models are provided in the SI appendix. The errors reported were standard errors to parameter fits obtained.

| **Langmuir One-Site Binding Model** |  |  |  |
| --- | --- | --- | --- |
|  |  | Cellulose I | Cellulose III |
| CBM1 | $n_{\text{max}}$ | $4.34 \pm 0.05$ | $3.32 \pm 0.09$ |
| | $K_d$ | $8.69 \pm 0.31$ | $10.55 \pm 0.91$ |
|  | RMSE | 0.17 | 0.17 |
| CBM3a | $n_{\text{max}}$ | $2.36 \pm 0.10$ | $1.48 \pm 0.11$ |
| | $K_d$ | $2.15 \pm 0.46$ | $6.15 \pm 1.75$ |
|  | RMSE | 0.22 | 0.17 |
| **Langmuir Two-Site Binding Model** |  |  |  |
|  |  | Cellulose I | Cellulose III |
| CBM1 | $n_{\text{max},1}$ | $4.14 \pm 0.01$ | $2.81 \pm 0.03$ |
| | $K_{d,1}$ | $13.68 \pm 0.13$ | $25.06 \pm 0.95$ |
| | $n_{\text{max},2}$ | $0.42 \pm 0.01$ | $0.75 \pm 0.03$ |
| | $K_{d,2}$ | $0.13 \pm 0.01$ | $0.92 \pm 0.07$ |
|  | RMSE | 0.11 | 0.12 |
| CBM3a | $n_{\text{max},1}$ | $1.81 \pm 0.03$ | $3.66 \pm 0.22$ |
| | $K_{d,1}$ | $16.64 \pm 1.35$ | $227.30 \pm 17.8$ |
| | $n_{\text{max},2}$ | $0.99 \pm 0.04$ | $0.67 \pm 0.02$ |
| | $K_{d,2}$ | $0.28 \pm 0.02$ | $0.64 \pm 0.05$ |
|  | RMSE | 0.17 | 0.12 |
| **Langmuir-Freundlich Binding Model** |  |  |  |
|  |  | Cellulose-I | Cellulose-III |
| CBM1 | $n_{\text{max}}$ | $4.80 \pm 0.02$ | $3.82 \pm 0.02$ |
| | $K_d$ | $6.90 \pm 0.04$ | $7.77 \pm 0.07$ |
| | $m$ | $0.77 \pm 0.00$ | $0.73 \pm 0.00$ |
|  | RMSE | 0.14 | 0.14 |
| CBM3a | $n_{\text{max}}$ | $3.21 \pm 0.05$ | $7.58 \pm 0.50$ |
| | $K_d$ | $2.61 \pm 0.06$ | $19.10 \pm 1.27$ |
| | $m$ | $0.53 \pm 0.01$ | $0.42 \pm 0.01$ |
|  | RMSE | 0.17 | 0.13 |

Since it was challenging to pick a suitable Langmuir model to identify a clear molecular-basis for reduced CBM1 or CBM3a binding to cellulose III, we also estimated the partition coefficients (i.e., *n*_*max*_*/K*_*d*_) for a larger library of several Type-A CBMs for both cellulose allomorphs at room temperature under non-saturating protein loadings (**Figure 2D** and raw data provided in SI Appendix **Fig. S7**). The relative binding order of CBM families based on estimated partition coefficient value was roughly similar between both allomorphs, where Family 1∼Family 2a << Family 64 < Family 3a < Family 5∼Family 10. However, the partition coefficients for all Type-A CBMs tested were always significantly lower for cellulose III. The decrease in partition coefficients ranged from 2-fold to 13-fold depending on the CBM family, with CBM64 and CBM1 showing the largest (∼13.3-fold) and smallest (∼2.3-fold) fold change between the two allomorphs, respectively. Note that the GFP domain alone had insignificant binding affinity towards either cellulose allomorph (∼0.1 L/g partition coefficient), suggesting that CBM domains alone were largely responsible for binding of GFP-CBM fusion proteins to either cellulose allomorph.

Although these results clearly show that Type-A CBMs have reduced binding to cellulose III under equilibrium conditions, measurement of adsorption and desorption rate constants can help further understand the rate limiting steps for native or pretreated cellulose hydrolysis by cellulases. Quartz crystal microbalance with dissipation (QCM-D) and fluorescence recovery after photobleaching (FRAP) based kinetic binding assays were also setup to show a 2 to 3-fold increased dissociation rate for CBM3a on cellulose III, as discussed in SI Appendix Results and Discussion section (see SI Appendix **Fig. S8-S9** and **Table S1**), that largely contributes to the reduction in bulk equilibrium binding affinity reported in **Table 1**. However, these equilibrium and kinetic binding assays do not provide information on the heterogeneity of binding interactions and possible binding orientations of CBMs on cellulose surfaces. We next developed a single-molecule CBM-cellulose binding assay to probe the multimodality of CBM-cellulose binding interactions various cellulosic allomorphs.

### Single-molecule CBM-cellulose bond rupture assay reveals multimodal nature of CBM binding

We designed an optical tweezers-based CBM-cellulose rupture assay to systematically characterize the binding interactions of a model CBM1 (from *T. reesei* Cel7A) towards Cladophora derived crystalline cellulose allomorphs. Our tweezer CBM-cellulose binding assay design is similar to our Cel7A motor motility assay reported previously (30). Here, GFP-CBM1 was tethered via a 1,010-bp DNA tether and attached to a 1.09 μm streptavidin-coated polystyrene bead (**Figure 3A**). Cellulose fibers were affixed to a glass coverslip.

**Figure 3.**
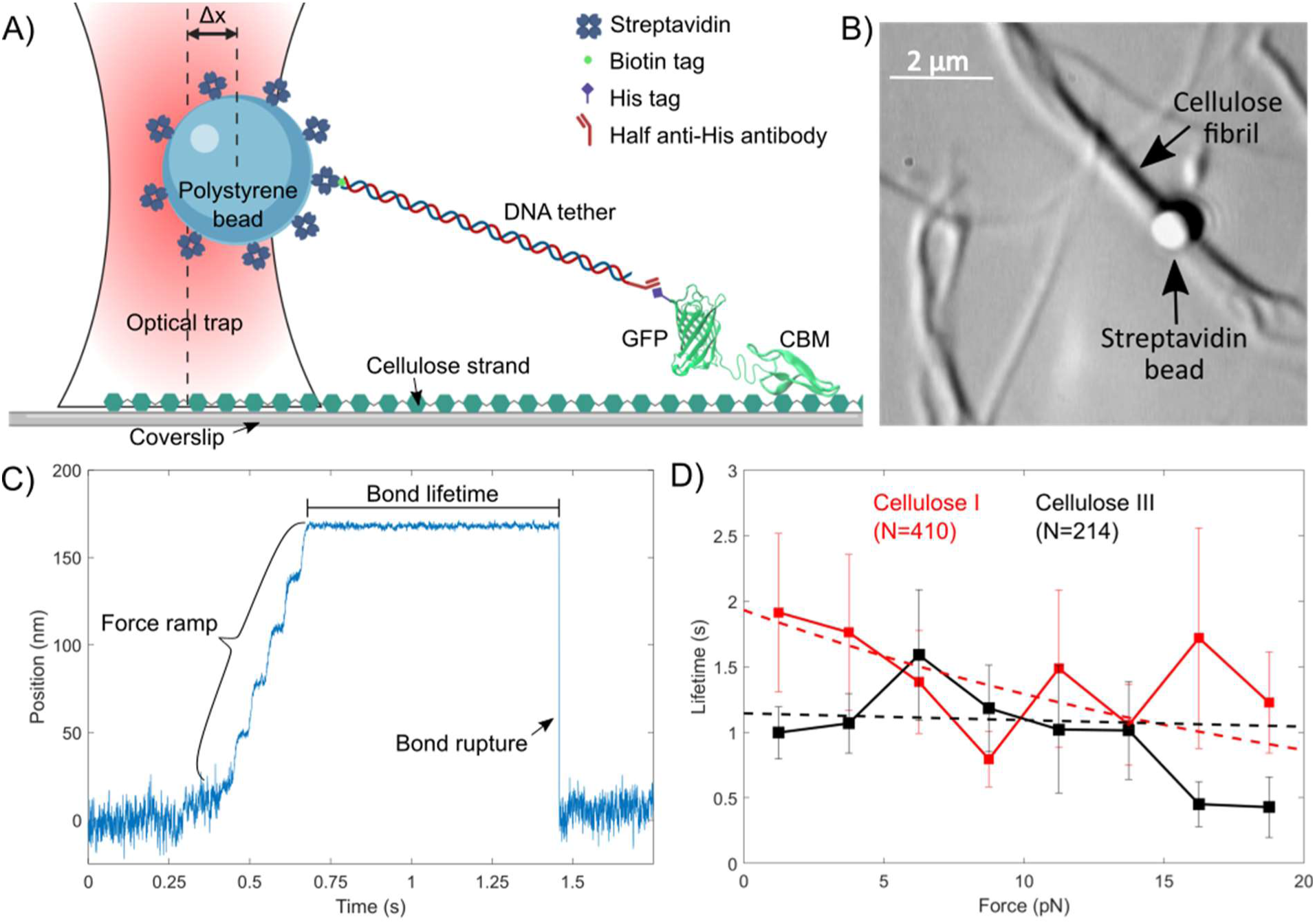
Optical tweezers based single-molecule ‘rupture’ assay reveals multimodal nature of CBM1-cellulose interactions. (A) Schematic of rupture assay setup (not to scale). A streptavidin coated bead is tethered to a single his-GFP-labeled CBM1 via a DNA linker containing an anti-His antibody Fab and a biotin tag on opposite ends. Δx represents the distance bead is displaced from trap center. Figure was created with BioRender.com. (B) Brightfield image of rupture assay showing Cladophora based cellulose microfibrils localized on the glass cover slip. CBM-cellulose binding is facilitated by moving the optically trapped bead close to the fiber. Bead position is tracked by a detection laser as force is loaded across the bond. (C) Representative position trace for a single CBM-cellulose rupture event showing bond lifetime and a single rupture is shown here. (D) Force vs Lifetime relationship for the CBM1-cellulose interaction on Cladophora cellulose I (black) and cellulose III (red) is shown. Lifetimes were binned at 2.5 pN intervals. Weighted single exponential fits are shown as dashed lines. Error bars depict standard error from the reported mean for each bin. N represents the total number of CBM-cellulose bond rupture events measured for each substrate. Additional supporting data can be found in SI appendix **Fig. S10-S12**.

For each assay run, individual beads were trapped and placed in the immediate vicinity of individual cellulose microfibers to facilitate a non-covalent CBM-cellulose bond (**Figure 3B**). Upon bond formation, the stage was moved to pull the DNA-tether taut and exert a force on the CBM-cellulose bond. Total bond lifetime and rupture force were then calculated for each individual CBM-cellulose interaction (**Figure 3C**). Hundreds of rupture events data from individual assay runs were pooled and binned at 2.5 pN intervals for cellulose I and cellulose III to generate force-lifetime distribution plots (**Figure 3D**). Raw force-lifetime scatterplots are provided in SI appendix **Fig. S10A**. Averaging all the rupture events, we find that the mean lifetime of CBM1 binding to cellulose I was 1.41 ± 0.20 s (SEM; N=410) and to cellulose III was 1.11 ± 0.12 s (SEM; N=214). The standard deviation of lifetimes of the CBM1-cellulose I and CBM1-cellulose III bonds were 4.12 s and 1.82 s, respectively. While one-way ANOVA test determined that the datasets over the entire rupture force range are not statistically different (**Fig. S10B**), slight differences can be seen at lowest (0-2.5 pN) and highest (17.5-20 pN) rupture force ranges. Furthermore, while the average lifetimes show different profiles, there was also a broad spread in the distribution of observed bond lifetimes with a great deal of overlap between cellulose I and cellulose III, indicating that multiple binding states with distinct characteristic bond lifetimes are possible for CBM1 binding to both cellulose I and III.

As seen previously for protein-ligand interactions in other single-molecule studies (37), CBM-cellulose binding was expected to show classic slip-bond behavior; i.e., as the rupture force increases, the total bond lifetime decreases. However, fits to the force-lifetime distribution failed to converge to a single exponential decay suggesting that multiple binding modes are likely present for CBM-cellulose. A classical unimodal slip bond would exhibit a single exponential decay (38), therefore it is clear that CBM1 does not follow this model when interacting with either cellulose allomorph. Binding of CBM1 on cellulose revealed a spread with a more complex multimodal and heterogenous binding behavior. In **Figure 3D**, weighted exponential decay fits (dashed lines) consistent with the expected slip-bond do not converge to conclusive unimodal behavior. This multimodal distribution was independent of the source of cellulose and similar results were also seen with filter paper cellulose (SI appendix **Fig. S11**). We also performed controls to test for artifacts associated with full anti-His antibody or Fab fragment binding but there was no major difference seen in the multimodal distribution of the force-lifetime results (SI appendix **Fig. S12**). Interestingly, the multimodal distribution of the force-lifetime was sensitive to the CBM structure as illustrated by the differences in rupture force-lifetime distribution seen for wild-type CBM1 and its Y31A mutant with a minor modification to the planar aromatic binding residue (SI appendix **Fig. S13**). Since it was difficult to directly determine an exact number of distinct CBM-cellulose binding states and their characteristic lifetimes using this rupture assay alone, we next conducted molecular dynamics (MD) simulations to provide an atomistic basis and better interpretation of the observed Cel7A tweezer motility and CBM1 binding results.

### CBM binding modalities are highly sensitive to cellulose surface ultrastructure

Based on the preferred binding orientation of CBM1 on model cellulose I and III crystal surfaces obtained from unbiased MD simulations (see **SI Appendix Fig. S14**), a potential of mean force (PMF) was calculated to estimate the CBM1 binding free energy during adsorption to the hydrophobic surface of distinct crystalline cellulose allomorphs. As shown in **Figure 4A**, in the case of cellulose I, only one PMF energy minimum well was observed corresponding to the dominant CBM1-cellulose configuration observed during the unbiased MD simulations whereby the Y31 residue faces the non-reducing end (i.e., canonical orientation based on native Cel7A favored activity from non-reducing end of cellulose). However, in the case of highly crystalline cellulose III, two PMF energy minima wells were observed, one in which Y31 faces the reducing end closer to the surface and another in which it faces the non-reducing end away from the surface. These configurations are annotated as non-canonical and canonical, respectively, in **Figure 4**. These two configurations are separated by roughly 0.2 nm in the PMF free energy diagram, where the distance is measured normal to cellulose surface, with a marginal energetic barrier of 2 kcal/mol separating the two minima basins. A closer examination of the CBM1 structure revealed that if the protein binds in the so-called ‘canonical’ orientation to the cellulose III surface at a shorter distance, then the Y5 residue exhibits significant steric clashes with the cellulose III adjacent surface chains (SI appendix **Fig. S14**). This explains why the ‘canonical’ CBM1 configuration is observed only at slightly longer distances away from the cellulose III surface. Irrespective of the preferred orientation for CBM1 to cellulose III surface and the degree of cellulose III crystallinity, the calculated free energy of binding for CBM1 was always lower for cellulose III compared to cellulose I. These results clearly support predictions from the bulk adsorption assays where the estimated equilibrium binding affinity for CBM1 was lower for cellulose III than cellulose I.

**Figure 4.**
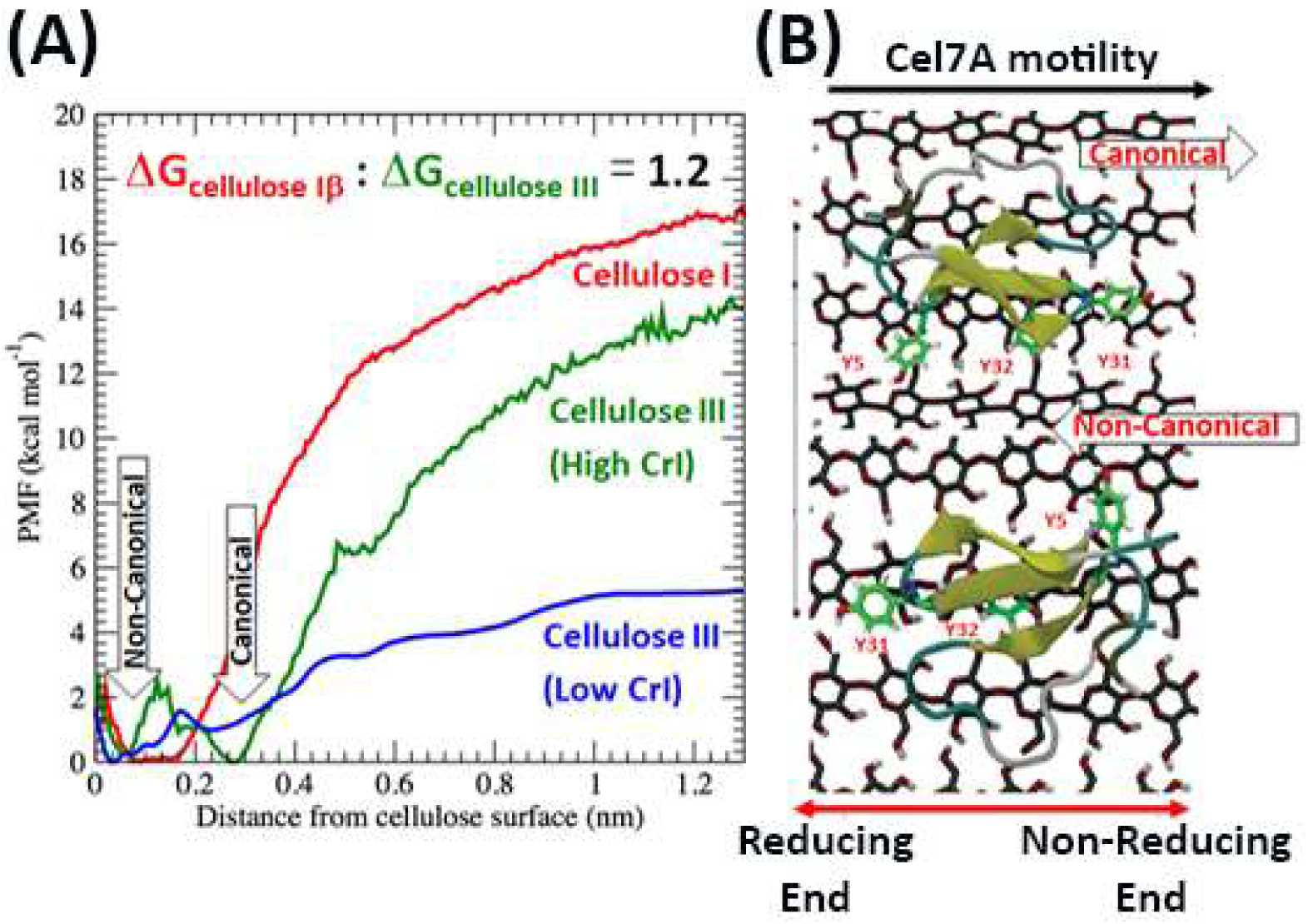
Molecular dynamics (MD) simulations provide an atomistic basis for distinct multimodal binding interactions of CBM1 to cellulose III allomorph surfaces. (A) Potential mean force calculations were carried out to estimate the binding free energy of CBM1 with cellulose allomorphs to show that binding free energy is 1.2-fold higher for cellulose I (in red) versus cellulose III. High crystallinity index (CrI; in green) and low CrI (in blue) models of cellulose III were studied here (see SI appendix for details). Note that the two energy wells for cellulose III correspond to the canonical and non-canonical orientations of CBM1. (B) Canonical orientation refers to Y5 residue facing the reducing end, as it favors the processive motility of Cel7A from reducing to non-reducing end of cellulose chain. Figure shows canonical (top) and non-canonical (bottom) orientations of CBM1 on high CrI cellulose III.

## Discussion

Carbohydrate-binding modules (CBMs) play an important role in polysaccharide degradation in nature, often by increasing the concentration of tethered catalytic domain to the targeted or associated adjacent polysaccharides (32, 39, 40). Mechanistic understanding of CBM-polysaccharide binding interactions also has implications for cost-effective industrial conversion of lignocellulosic biomass to fuels and chemicals by cellulase-mediated hydrolysis (41, 42). However, we still lack a complete understanding of the role of CBMs in processive motility of industrial workhorse enzymes such as Cel7A. Although molecular simulations have been employed to study processive cellulase cycle steps such as chain decrystallization (43), glycosylation (22), deglycosylation (23) and dissociation (44), the role of CBMs in initial motility commitment of catalytic domains is not yet clear. Moreover, lignocellulosic biomass often undergoes pretreatment in a biorefinery, which can sometimes lead to modification of cellulose crystal structure to form non-native structures such as cellulose III (11, 14). From an evolutionary standpoint, Type-A family CBMs and cellulase catalytic domains have naturally evolved to breakdown native plant-based cellulose I allomorph (45). Therefore, there is a need to better understand the complex binding interactions of CBMs to both native allomorphs such as cellulose I and non-native allomorphs such as cellulose III using multifaceted characterization tools to gain a truly molecular-level understanding of CAZyme-polysaccharide interactions.

Here, we have used classical pull-down binding assays, single molecule optical tweezer-based bond rupture assays, and molecular dynamics simulations to obtain a comprehensive understanding of CBM-cellulose binding interactions with a focus on cellulose I and III allomorphs. Classical pull-down binding assays (also called solid-state depletion assays) have been employed extensively to study protein binding to insoluble polysaccharides like cellulose (8). Under the assumption that conditions of Langmuir adsorption have been met (46), simple mathematical models are often used to extract physicochemical parameters such as bulk binding affinities and number of available sites for CBM binding to cellulose (7). Researchers decide between the choice of multi-site Langmuir models, based on minor differences in the overall model fit, or an *ad hoc* basis with no clear justification and the limitations of such approaches have been outlined by Igarashi and others (46–48). Our model-fitting analysis for CBM1 binding, revealed a 1.3-fold reduction in binding affinity for cellulose III using one-site binding model which magnified to ∼2 to 5-fold difference when using two-site binding models. Hence, interpretation of classical binding assay results could vary drastically depending on the exact model used. This issue is further exacerbated by the presence of fewer data points in the higher free protein concentration range going up only to 15-25 μM (47, 49), as opposed to a maximum of ∼250 μM tested for CBM1 in our case. Hence, we also performed truncation analysis by trimming down our binding data set for CBM1 to exclude higher protein concentrations (i.e., included maximum concentrations <15 μM or <50 μM). We observed that the number of predicted binding sites for both cellulose I and cellulose III decreased by ∼1.3 to 1.8-fold for the truncated datasets (see SI Appendix **Table S2**). Interestingly, our truncated dataset fitted models predicted a slightly weaker affinity of CBM1 towards cellulose I versus cellulose III, which was contrary to predictions made from full data set. In addition, Langmuir adsorption models can only be applied in the absence of protein denaturation and protein-protein interactions at the binding interface (46). Here, we also studied the binding of a small-molecule CBM-surrogate calcofluor which does not have any associated issues such as denaturation seen for proteins (see **SI Appendix Fig. S15**). However, we also observed concave upward Scatchard plots for calcofluor binding to cellulose, indicating the presence of multiple overlapping binding sites. Overall, analysis of our classical pull-down binding assay data highlights the critical limitations of using multi-site Langmuir binding models for studying protein adsorption to heterogeneous substrates such as cellulose, without a molecular basis and the misleading interpretations that can result from such analyses.

Single-molecule force spectroscopy has been employed previously to distinguish the nature of protein-ligand bonds (37) and hence infer multi-modality or conformational transitions involved in protein-ligand binding interactions (50). Single-molecule force spectroscopy in conjunction with steered MD simulations has been employed extensively to study superior mechanical stability (51), characterize the force-unfolding behavior (52), and resolve multiple binding modes of cohesin-dockerin complexes (53). However, the application of AFM-based force spectroscopy to study CBM-cellulose binding has revealed challenges in distinguishing specific vs non-specific interactions (54). Here, we have developed a novel single-molecule optical tweezer-based bond rupture assay with piconewton (pN) force resolution and millisecond (ms) time resolution (50), to understand the heterogeneity of CBM binding behavior and provide a firm molecular basis for using a certain adsorption model to interpret classical ‘pull-down’ assay data. CBM1 showed multi-modal force-lifetime behavior towards both cellulose I and cellulose III, as evidenced by the inability to fit this dataset to either single or double exponential decay functions. We speculate that this multimodal behavior indicates multiple classes of overlapping binding sites with contributions from different classifications of cellulose substructures (35) namely crystalline regions with varying degrees of disorder, different crystal binding faces (55), and varied binding orientation/modes of CBM binding on the hydrophobic face of crystalline cellulose. However, due to the highly crystalline nature of our Cladophora derived cellulosic substrates (∼95%) and the observation that CBM1 binds predominantly to one crystal face (55), we speculate that the resulting multi-modality could largely arise from multiple equilibrium binding modes of CBM1 on preferred cellulose surfaces. This can be visualized from a purely geometric standpoint based on our simple CBM binding orientation model inspired from the original Buffon needle problem (56). Our model shows that the geometric distribution of CBM1 binding orientation states mostly align along the cellulose chain axis versus across the the chain axis under the key assumption that these states are energetically equivalent, as discussed in the SI Appendix Results and Discussion section (see **SI Appendix Fig. S16**). Future work combining site-directed mutagenesis of CBMs, force spectroscopy rupture assays, and MD simulations is necessary to test the impact of specific CBM binding motif mutations on altering certain binding modalities as illustrated by Jobst *et al*. (53). In relation to the classical pull-down binding assay results for CBM1, our single-molecule CBM-cellulose bond rupture assay reveals that the binding behavior cannot be explained by presence of just one or two classes of unique binding sites. However, fitting high-quality binding assay data to a Langmuir one-site model can still yield an average affinity arising from a combination of binding sites or modes, rather than data overfitting via a two-site model. In summary, our analysis recommends use of Langmuir one-site model to obtain binding parameters while also using complementary approaches to characterize the molecular-level origins of different binding modalities.

Molecular dynamics simulations have been employed extensively to study cellulolytic enzymes (22, 57–60), and can offer detailed atomistic insights into the highly heterogeneous and multimodal CAZyme binding interactions with insoluble polysaccharides. MD simulations have also revealed structural and dynamical features of cellulose III such as hydrogen bonding patterns, solvent accessible surface area, and decrystallization free energy (17, 43). While few studies have been carried out to understand CBM binding to cellulose, most work has been restricted to native cellulose I allomorph (61–63). Beckham-Nimlos-Alekozai have made several important observations concerning CBM1 binding to cellulose I, namely: (a) CBM1 binds predominantly to the hydrophobic cellulose face as opposed to the hydrophilic face with a ∼2.5 kcal/mol difference in binding free energy, (b) the planar aromatic motif of CBM1 binds preferentially to the hydrophobic face of cellulose I with thermodynamic stability along the cellulose chain axis corresponding to every repeating cellobiosyl unit, and (c) CBM1 prefers to bind such that the Y31 residue mostly faces the non-reducing end of cellulose I and this ‘canonical’ binding preference has likely co-evolved based on the appended Cel7A CD catalytic activity preference (61–63). In the current study, we only considered CBM1 binding to the hydrophobic surfaces for both cellulose I and III allomorphs. Our results mostly corroborate previous findings for CBM1 binding to cellulose I. Nevertheless, our unbiased MD simulation results (see **SI Appendix Fig. S14**) indicated that CBM1 has increased diffusivity on cellulose III surface and the planar aromatic residues exhibited increased root mean square fluctuation (RMSF) as compared to cellulose I. The increased RMSF of planar aromatic residues (Y5, Y31, Y32) on cellulose III arises due to improper stacking interactions with cellulose sheet, possibly due to the stepped nature of adjacent cellulose III surface chains that is not complementary to the natively evolved planar CBM1 binding motif. These results also support our observation of lower binding affinity of CBM1 toward cellulose III in classical binding assays. In addition, the possibility of CBM1 binding to cellulose III in the ‘non-canonical’ orientation can partially explain the difficulty in initial motility commitment for Cel7A on cellulose III based on our motility data. CBM1 plays an oft-neglected synergistic role in the association of Cel7A catalytic domain to cellulose that likely fine-tunes the subtle balance between productive versus non-productive binding (64). Our motility assays have, for the first time in reported literature, captured this early step of catalytic domain complexation to a reducing end before catalytic processive cycle begins. Future work will address the role of CBMs in both the association process and dissociation process, to obtain a greater understanding of the relationship between binding affinity and overall catalytic efficiency for processive cellulases (21).

## Materials and Methods

See SI appendix (Supplementary Text) for all materials and methods relevant to this study.

## Acknowledgments

The authors acknowledge support from the NSF CBET awards (1604421 and 1846797), ORAU Ralph E. Powe Award, NSF MCB award (1330792), NIH Grant (R01GM101001), Rutgers Global Grant, Rutgers Division of Continuing Studies, Rutgers School of Engineering, DOE Bioimaging Award (Office of Science DE-SC0019313) and the Great Lakes Bioenergy Research Center (DOE BER Office of Science DE-FC02-07ER64494). SPSC would like to particularly thank Professor Brian Fox (UW Madison) and Professor Bruce Dale (MSU) for kindly providing access to their lab’s resources at the onset of this project for generation of relevant plasmid DNA and cellulose substrates. Special thanks to Izak Smith for help with collection of Cladophora algae from the Yahara River-Lake Watershed, Sungsoo & Amy Lim for help with generation of GFP-CBM constructs, Leonardo Sousa for conducting ammonia pretreatment, Shashwat Gupta for XRD data analysis, Ki-Bum Lee & Hyeon-Yeol Cho for access to the AFM, and Umesh Agarwal for access to the FT-Raman spectrometer. BN and JMY would like to thank Dr. Ashutosh Mittal (NREL) for his help with XRD measurements of Avicel cellulose derived nanocrystals. BN would also like to thank Dr. Jeff Linger at NREL for providing access to lab resources to perform QCM-D experiments.

