## Supplementary material for "Multiscale Characterization of Complex Binding Interactions of Cellulolytic Enzymes Highlights Limitations of Classical Approaches": SI Appendix


**This PDF file includes:**

- Supplementary text (SI Materials, Methods, and Results)
- Figures S1 to S16
- Tables S1 to S3
- Legends for Movies S1 to S3
- SI References

**Other supplementary materials for this manuscript include the following:**

- Movies S1 to S3

**SI Appendix Materials and Methods**

***Crystalline Cellulose Isolation and Anhydrous Liquid Ammonia Pretreatment:*** High crystallinity cellulose I (called native Cladophora cellulose I) from *Cladophora* sp. (*Cladophora glomerata*) was isolated and characterized as described previously (1). High purity (>98% cellulose content, dry weight mass basis or dwb) plant-derived microcrystalline cellulose I (called Avicel cellulose I) was purchased from Sigma-Aldrich (Avicel PH-101, Lot No. BCBD6923V). These native cellulose samples were used to generate respective Avicel or Cladophora derived crystalline cellulose III using a suitable anhydrous liquid ammonia based pretreatment process (2, 3). All cellulose III samples were kindly generated by Dr. Leonardo Sousa using a general protocol highlighted elsewhere (3). Briefly, cellulose III was prepared in a high-pressure stirred batch reactor at 90 °C for 30 min (for Avicel) or 4 h (for Cladophora) residence time using at least a minimum 6:1 ammonia-to-cellulose loading ratio (dwb). The reactor pressure was maintained constant at 1000 psi using nitrogen gas during the pretreatment, and ammonia was slowly evaporated from the reactor through a venting valve after the desired residence time. During this evaporation process, the temperature of the reactor was slowly decreased and kept stabilized at 25 °C. The treated cellulose sample was then removed from the reactor and placed overnight in the fume hood to evaporate any residual ammonia. All treated cellulose samples were stored at 4 °C in a zip sealed bag prior and were used directly without any further drying.

***Cellulose Characterization using XRD & FT-Raman Spectroscopy:*** Details regarding the X-ray diffraction (XRD) method and data analysis methods/results are provided elsewhere (2, 3). Briefly, XRD was performed on an X-ray diffractometer with beam parallelized by a Gobel mirror (D8 Advance with Lynxeye detector; Bruker, Bruker AXS Inc., Madison, WI, USA). CuKα radiation (wavelength = 1.5418 Å) was generated at 40 kV with 40 mA current and the detector slit was set to 2.000 mm. Samples were analyzed using a coupled 2θ/θ scan type with a continuous PSD fast scan mode. The 2θ started at 8.000° and ended at 30.0277° with increments of 0.02151°, while θ started at 4.0000° and ended at 15.0138° with increments of 0.01075°. Step time was 1.000 sec (i.e., 1025 total steps, effective total time 1157 sec per run). Dry cellulose samples (approximately 0.5 g) were placed in a specimen holder ring made of PMMA with 25 mm diameter and 8.5 mm height, rotating at 5 degrees per minute during analysis. Cellulose crystallinity was estimated based on the Segal peak height and amorphous peak deconvolution based methods (4, 5). Please note that Miller indices used in this paper for each contributing predominant diffraction peak/s conform to the convention with 'c' as the fiber axis, a right-handed relationship among the axes and the length of a<b, as recommended recently by Alfred French (6), to avoid confusion with other naming conventions. Briefly, for the XRD Segal peak height method, cellulose crystallinity index was calculated from the ratio of the height of the (110) or (200) plane equatorial reflection peak and the height of the minimum between the (110) or (200) and (010) or (110) plane equatorial reflection peaks for Cladophora or Avicel PH-101 cellulose I, respectively. For cellulose III, cellulose crystallinity index was calculated from the ratio of the height of the (100) plane equatorial reflection peak and the height of the minimum between the (100) and (002) plane equatorial reflection peaks. Note that, the three main peaks for native Cladophora cellulose I one-chain triclinic unit cell have Miller indices of (100), (010) and (110), which are the counterparts to the (1-10), (110) and (200) peaks of Avicel PH101 cellulose I pattern. Peak deconvolution methods have been used extensively to calculate cellulose crystallinity index (5, 7–9). XRD peak deconvolutions were carried out using PeakFIT (Version 4.12, Systat Software Inc, San Jose, CA) as described elsewhere (2, 5). For all peak deconvolutions F values are always > 30,000 while R-squares > 0.999. Similar to previous work (3, 8, 10, 11), XRD equatorial reflections for (100), (010), and (110) crystallographic planes for native Cladophora cellulose I were at approximately 14.9°, 17.1°, and 23.0° Bragg angles (2θ), respectively. Similar to previous work (3, 12), equatorial reflections for (010), (002), and (100) crystallographic planes for Cladophora cellulose III were at approximately 11.8°, 17.4°, and 20.9° Bragg angles (2θ), respectively. While, XRD equatorial reflections for (1-10), (110), and (200) crystallographic planes for Avicel PH-101 based cellulose I were at approximately 14.9°, 16.3°, and 22.5° Bragg angles (2θ), respectively. And equatorial reflections for (010), (002), and (100) crystallographic planes for Avicel PH-101 based cellulose III were at approximately 11.7°, 17.1°, and 20.6° Bragg angles (2θ), respectively. Avicel derived cellulose allomorphs crystallinity index calculated using these methods are also reported in a recent paper (3), while Cladophora crystallinity index is reported here in SI appendix results section.

Additional supporting details regarding the FT (Fourier Transform) Raman based spectroscopic characterization methods/results are provided as well. Briefly, a MultiRam FT-Raman spectrometer (Bruker) was used to collect Raman spectra for cellulose samples. The FT-Raman spectrometer was equipped with a 1064-nm 1000-mW Nd:YAG laser. For Raman analysis, cellulose pellets were first prepared from either air-dried or lyophilized samples prior to analysis. In most cases, spectra with high signal-to-noise (S/N) ratios was obtained upon using a 660 mW laser power setting and collecting over 512 scans per sample. The spectra were converted to ASCII format and exported to Microsoft Excel for direct plotting/analysis. The interconversion of cellulose I to III was confirmed based on previously published reports using Cladophora or cotton linters derived cellulose allomorphs (3, 13–16). Peak assignments of the vibrational spectrum of cellulose I and III have been described elsewhere (3, 15, 17). Briefly, 250-550 $cm^{-1}$ region for cellulose has predominant group motions attributed to skeletal-bending modes involving C-C-C, C-O-C, O-C-C, and O-C-O internal bond coordinates. The 550-750 $cm^{-1}$ region corresponds to mostly out-of-plane bending modes involving C-C-C, C-O-C, O-C-O, C-C-O, and O-H internal bond coordinates. The peaks around 900 $cm^{-1}$ are shown to involve bending of H-C-C and H-C-O bonds localized at C-6 atoms of the hydroxymethyl group. The 950-1200 $cm^{-1}$ region corresponds to mostly stretching motions involving C-C and C-O internal bond coordinates. The 1200-1500 $cm^{-1}$ region corresponds to mostly bending motions involving H-C-C, H-C-O, H-C-H, and C-O-H internal bond coordinates. The region of 1400-1500 $cm^{-1}$ for cellulose has been shown to be particularly sensitive to the $CH_2$ scissor bending modes that are sensitive to the Trans-Gauche or TG (1480 $cm^{-1}$) and Gauche-Trans or GT (1460 $cm^{-1}$) conformations of the hydroxymethyl group (15).

***CBMs gene synthesis and cloning:*** *E. coli* codon optimized genes encoding CBMs, with additional flanking *AflII* and *BamHI* restriction sites, inserted into a standard pUC57-Kan vector were ordered from Genscript USA Inc (Piscataway, NJ). DNA sequences for all CBMs are provided in the table below. An *E. coli* expression vector pEC-GFP-CBM3a was kindly provided by the Fox lab (UW Madison). Sequence information regarding the family 3a CBM from *Clostridium thermocellum* expressed using this pEC vector have been published already (18). The pEC vector sequence map and strategies for primer design and CBM genes sub-cloning have been reported already (18, 19). Briefly, polymerase incomplete primer extension (PIPE) based ligation independent cloning approach was used to transfer the CBM nucleotide gene sequences from the respective pUC57 to pEC vector (20). For the creation of CBM1 Y31A mutant used for bond rupture assay (reported in **Fig. S13**), site-directed mutagenesis with complementary forward and reverse primers was used. Polymerase chain reaction (PCR) was catalyzed by Herculase II Fusion DNA polymerase (Agilent Technologies, Santa Clara, CA). Destination pEC vector and CBM insert gene amplification was carried out using suitably designed PIPE primer pairs. The PCR amplification of pUC57 and pEC vectors using corresponding vector/insert PIPE reactions primer pairs were carried out in separate tubes. After PCR, respective CBM PIPE reaction product aliquots (2 µL) were mixed together and immediately transformed into competent *E. coli* E. cloni 10G cells (Lucigen, Madison, WI). If the PIPE cloning strategy was not successful, the pUC57 and pEC vectors were digested using *AflII* (New England Biolabs, Ipswich, MA) and *BamHI* (Promega, Madison, WI) restriction enzymes. The restriction enzyme products were ligated using T4 DNA ligase (New England Biolabs) and the ligation mixture was instead transformed into competent *E. cloni* 10G cells. Individual transformant colonies were next screened by PCR amplification and transformants containing inserts with the approximate correct size were identified by agarose gel electrophoresis. Plasmids isolated from positive colonies were sequenced to confirm nucleotide identity at the UW Biotechnology Center (and/or at Genscript, Piscataway, NJ). Transformed strains were stored as 20% glycerol stocks were maintained at -80 ˚C, while all relevant pEC-GFP-CBM plasmids were also maintained at -80 ˚C for long-term storage. The sequence-verified plasmid, named pEC-GFP-CBM, encoded a 5' $His_8$-tag, followed by *gfp*, a linker sequence, and the relevant *cbm*. The linker sequence encoded for a 42 amino acid linker peptide reported previously by Takasuka and co-workers (21). Here, we have characterized in detail two distinct but structurally homologous family 2a CBMs from *Streptomyces* sp. SirexAA-E and *Acidothermus cellulolyticus*. *Streptomyces* sp. SirexAA-E was recently identified by GLBRC researchers for its high cellulolytic ability (22). In addition, we have also characterized two distinct but structurally homologous family 1 CBMs from *Trichoderma reesei* belonging to Cel7A (cellobiohydrolase I or CBHI) and Cel6A (cellobiohydrolase II or CBHII) processive cellulases. However, here we exclusively report results for CBM1 from Cel7A, unless mentioned otherwise. Previous studies have shown that the structure and function relationship of homologous CBMs belonging to the same family are often similar, as also seen in our current work when comparing binding results between the two family 2a CBMs.

| CBM Family | CBM Gene Sequence | Organism Source |
|---|---|---|
| 1 | CCGGGTCCGACCCAGAGCCATTATGGCCAGTGCGGTGGTAT TGGTTATAGCGGTCCGACCGTGTGCGCAAGCGGTACCACCT GCCAGGTGCTGAACCCGTATTATAGCCAGTGCCTG | *Trichoderma reesei* |
| 2a | GCGGCGAGCGGCGCACGTTGCACCGCAAGTTATCAAGTGAA TAGCGATTGGGGGAACGGTTTCACGGTTACCGTCGCAGTTA CCAACTCAGGTTCTGTTGCTACCAAAACCTGGACGGTGTCGT GGACCTTCGGCGGTAATCAGACTATCACCAACAGCTGGAAC GCGGCGGTCACACAGAACGGCCAGAGTGTGACTGCACGTA ACATGAGCTACAATAATGTTATTCAACCAGGCCAAAATACGA CCTTTGGTTTTCAAGCCTCGTACACGGGCAGTAACGCAGCA CCGACCGTTGCGTGCGCGGCGAGT | *Acidothermus cellulolyticus* |
| 2a | Refer to Lim et al. 2014 (19) | *Streptomyces sp. SirexAA-E* |
| 3a | Refer to Whitehead et al. 2017 (18) | *Clostridium thermocellum* |
| 5 | ATGGGTGATTGTGCTAACGCAAATGTCTATCCGAACTGGGTG TCTAAAGATTGGGCGGGTGGTCAACCGACGCATAACGAAGC GGGTCAGAGCATTGTGTATAAAGGCAACCTGTACACCGCGA ATTGGTACACCGCATCAGTGCCGGGTTCAGACTCATCGTGG ACGCAGGTTGGTAGTTGTAATTGA | *Erwinia chrysanthemi* |
| 10 | ATGGGCAATCAACAATGTAACTGGTATGGCACCCTGTATCCG CTGTGTGTGACGACGACGAATGGCTGGGGCTGGGAAGATCA ACGCAGCTGCATCGCCCGTAGCACCTGCGCGGCTCAACCG GCACCGTTTGGCATCGTGGGTAGCGGCTGA | *Cellvibrio japonicus* |
| 64 | CCGACCCCGTCTGGCGAATATACGGCGATTGCCCTGCCGTT TACCTACGATGGCGCCGGTGAATATTACTGGAAAACCGACC AATTCAGCACCGATCCGAATGACTGGTCACGTTATGTCAACT CGTGGAATCTGGATCTGCTGGAAATTAACGGTACCGACTACA CGAATGTGTGGGTTGCACAGCATCAAATCACGCCGGCTAGT GATGGCTACTGGTATATTCACTACAAAGGCTCGTATCCGTGG TCGCATGTGGAAATCAAA | *Spirochaeta thermophila DSM 6192* |

***Small-scale expression testing of GFP-CBMs:*** pEC-GFP-CBM plasmids were first transformed into BL21-CodonPlus-RIPL [λDE3] (Stratagene, Santa Clara, CA) or Rosettagami 2 [DE3] (Novagen, Santa Clara, CA) *E. coli* competent strains for small-scale protein induction/expression optimization screening. After cells were grown to the optical density of 0.5-0.7 (mid-exponential phase) in a non-inducing medium (23), expression was induced in either an auto-induction medium (23) at 25 ˚C or in LB medium with varying concentrations of IPTG (0.1-1mM) during incubation at 16 ˚C, 25 ˚C, or 37 ˚C. Apart from GFP-CBM2a (ActE), as also reported previously (19), soluble cytoplasmic protein expression was observed for all other GFP-CBMs in nearly all of the expression conditions tested. Superfolder enhanced GFP (or eGFP) tag has been reported to increase soluble fusion protein expression yields by likely preventing aggregation of hydrophobic proteins like CBMs (24). Nevertheless, small-scale immobilized metal affinity chromatography with $Ni^{2+}$-NTA (immobilized metal affinity chromatography or IMAC) purification and cellulose binding assays were conducted for all soluble protein fractions to confirm cellulose-binding activities and hence identify optimum cell culture conditions for large-scale protein production/purification to run all reported bulk and single-molecule CBM binding assays.

***Large-scale expression of GFP-CBMs:*** *E. coli* BL21-CodonPlus-RIPL [λDE3] (Stratagene, Santa Clara, CA) or Rosettagami 2 [DE3] (Novagen, Santa Clara, CA) competent strains were transformed with the relevant pEC-GFP-CBM plasmid based on the small-scale expression results. Suitable transformants were inoculated into 50 mL of chemically defined non-inducing medium (23), in the presence of 50 µg/mL kanamycin and 25 µg/mL chloramphenicol selection antibiotics. The non-inducing medium contained 2 mM $MgSO_4$, a 1:1000 dilution of trace metal salts mixture (equivalent to 50 mM $Fe^{3+}$, 20 mM $Ca^{2+}$, 10 mM $Mn^{2+}$, 10 mM $Zn^{2+}$, 2 mM $Co^{2+}$, 2 mM $Cu^{2+}$, 2 mM $Ni^{2+}$, 2 mM $Mo^{6+}$, 2 mM $Se^{4+}$, 2 mM $H_3BO_3$) into the medium, 0.5% glucose, 0.25% aspartate, 50 mM $NH_4Cl$, 25 mM $KH_2PO_4$, 25 mM $Na_2HPO_4$, 5 mM $Na_2SO_4$, 0.01% methionine, 1% of 17 amino acids (except cysteine, tyrosine, and methionine) each, and a vitamin cocktail (200 nM of vitamin B12, nicotinic acid, pyridoxine, thiamine, p-aminobenzoic acid, and pantothenate; 5 nM folic acid, and riboflavin). The culture was incubated overnight at 25 ˚C and then used to inoculate 2 liters of auto-induction medium (23). The auto-induction medium contained 1.2% tryptone, 2.4% yeast extract, 2.3% $KH_2PO_4$, 12.5% $K_2HPO_4$, 0.375% aspartate, 2 mM MgSO4, 0.8% glycerol, 0.015% glucose, and 0.5% α-lactose. The cultures were grown at 25 ˚C for ~27 h. The cells were harvested by centrifugation and the cell pellet was stored at -80 ˚C until further use. All chemicals were purchased from Sigma-Aldrich (St. Louis, MO).

***Purification of GFP-CBMs:*** The recovered cell pellet was thawed and re-suspended in 150 mL of ice cold 20 mM phosphate, pH 7.4, containing 500 mM NaCl, 20% v/v glycerol, 10 µg/ml lysozyme, and a protease inhibitor cocktail (containing benzamidine, EDTA and E-64 protease inhibitor from Sigma-Aldrich). The cells were sonicated with an ultrasound sonicator (550 Sonic Dismembrator, Fisher Scientific, Pittsburgh, PA) fitted with a microprobe (1-inch probe diameter) at 4 ˚C for 5 min with 10-s on-bursts and 30-s off periods. The cell debris containing the inclusion bodies was pelleted at 21,000 rpm at 4 ˚C (30 min) and the supernatant was collected in all cases except for GFP-CBM2a (ActE). Details regarding GFP-CBM2a (ActE) expression and purification are provided elsewhere (19). Briefly, due to the insolubility of the expressed GFP-CBM2a (ActE) under all conditions tested, this protein construct was first isolated from inclusion bodies, refolded, and then purified using IMAC as described previously (19). For all other GFP-CBMs, IMAC using $Ni^{2+}$-NTA based columns/media (GE Healthcare) was first used to isolate and purify $His_8$-tagged proteins from the *E. coli* cell lysate. All column-based protein purifications were carried out on a ÄKTA-FPLC system (GE Healthcare, Pittsburgh, PA). The cell lysate supernatant was first loaded onto the IMAC column at a medium flow rate of 1 – 2 ml/min. The column was then washed with buffer A (100 mM MOPS, pH 7.4, containing 10 mM imidazole and 100 mM NaCl), followed by additional washing using 95% IMAC buffer A spiked with 5% IMAC buffer B (100 mM MOPS, pH 7.4, containing 500 mM imidazole and 100 mM NaCl), and last followed by elution in 100% IMAC buffer B at a flow rate of 5 ml/min. Protein purity and molecular weight at each stage of the protein purification process was examined by SDS-PAGE (Criterion XT Bis-Tris Precast Gels, Bio-Rad). The presence of partially cleaved GFP-CBMs was identified in the IMAC-B eluents for some protein constructs (namely CBM1, CBM2a, CBM5, CBM10), which necessitated further purification using an amorphous cellulose or hydrophobic interaction affinity-based purification method, as already outlined elsewhere (19, 25), to isolate the intact protein fractions. Briefly, for cellulose affinity-based purification method, IMAC-B protein eluents were directly applied to a phosphoric acid swollen amorphous cellulose (PASC) media at the recommended loading (~200 mg crude protein added per gram dry weight cellulose) for preparative-scale purification (25). The amorphous cellulose slurry was prepared ahead of time and preequilibrated in a 50 mM pH 6.5 MES buffer (equilibration buffer or buffer A) at the desired solids concentration (10 g/L), prior to addition of the IMAC-B protein eluent. The crude protein-cellulose slurry was then intermittently and gently mixed at room temperature for a total incubation time of 0.5 h. The protein bound to PASC was then separated from the unbound protein in the supernatant by gentle centrifugation at 3500-g for 10 min. The recovered PASC pellet was then resuspended in a wash buffer (i.e., equilibration buffer+1M NaCl), using a 4:1 buffer to PASC pellet ratio (v/v), and gently mixed at room temperature for 10 mins to remove non-specifically bound proteins. The recovered PASC pellet containing the adsorbed GFP-CBMs was then finally suspended in 100% ethylene glycol elution solution, using a 4:1 glycol to PASC pellet ratio (v/v). The final ethylene glycol concentration of ~80% (v/v) was sufficient to elute a significant fraction of reversibly bound GFP-CBMs into the supernatant. The eluted protein rich supernatant was separated from PASC pellet and stored in 80% glycol solution at −20 ˚C for short term storage or immediately concentrated using IMAC columns prior to buffer exchange into 10 mM pH 6.5 MES (or pH 5.5) buffer and storage at −80 ˚C for long term storage in 0.5-1 ml aliquots. The molecular weight of the intact purified GFP-CBM monomers was confirmed by SDS-PAGE to match with the predicted translation products. Protein concentrations were estimated spectrophotometrically at 280 nm using the extinction coefficients calculated from the amino acid sequences for each construct. The histidine tags were not removed and have been reported to not influence CBM binding to cellulose (19, 26).

***GFP-CBM or calcofluor dye 'pull-down' binding assays with cellulose allomorphs:*** Cladophora based cellulose allomorphs, prepared as described above, were used to estimate binding affinity and partition coefficients for GFP-CBMs. All cellulose binding assays were carried out in 2 mL microcentrifuge tubes using 5 mg of pre-weighed cellulose suspended in 500 μL of 10 mM MES buffer, at pH 5.5 (for CBM1) or 6.5 (for all other CBMs), containing 2.5 mg/mL of bovine serum albumin (BSA) to minimize non-productive protein binding to the tube wall. Purified GFP-CBMs aliquots were thawed and buffer-exchanged (PD-10 desalting column, GE Healthcare) into 10 mM MES, at desired pH, and added to each 2 mL tube to achieve a final solution concentration. The final concentration typically ranged from 1-500 μg/mL for estimating partition coefficients or from 1 μg/mL to 10 mg/mL for full scale binding affinity measurements. The tubes were mixed in an orbital mixer set at 1000 rpm (ThermoMixer, Eppendorf) at 25˚C for 1.5-2 h to allow binding equilibrium to be reached. Controls with only proteins but no substrate were mixed to track total added protein concentration, while never-mixed controls with only proteins/buffer were used as controls to account for possible protein loss during mixing. Insoluble cellulose was recovered by centrifugation at 13,000 rpm for 2 min and 200 μL of the supernatant (w/wo suitable dilution in identical buffer) containing unbound protein was assayed for GFP fluorescence (488 nm excitation and 509 nm emission; with 495 nm cut-off) to quantify the total fraction of bound protein versus the original added amount (2). All assays were performed in at least duplicate for each protein loading condition (~22 protein loadings). Error bars reported for all bulk binding assays represent one standard deviation (±1σ) from mean values from duplicates.

Calcofluor White dye (Sigma-Aldrich) binding assays on Avicel PH-101 based cellulose I and III were performed in 1.5 ml Eppendorf tubes at varying dye concentrations, similar to previously reported cellulose-solute binding assay methods (2, 27). Briefly, 5 mg of Avicel cellulose-I or cellulose-III was added to each well by pipetting suitable aliquots of a uniformly suspended cellulose slurry prepared in deionized water. Suitable calcofluor dilutions were prepared in deionized water (at pH 7.0) to achieve the desired concentration in each tube. Note that 30 mM NaCl was added to each tube. For cellulose blanks, deionized water alone instead of cellulose was added to make up the volume instead to the final desired 620 μl level. All binding assays were performed with three replicates per assay condition. The tubes were then shaken in Thermomixer at 1000 rpm for 1 h at room temperature. After 1 hour, all tubes were centrifuged at 13,000 rpm for 5 minutes and supernatant was removed from each well to be transferred into opaque microplates for reading calcofluor fluorescence. The plates were read using a Molecular Devices M5e spectrophotometer at the following settings: 365 nm excitation, 450 nm emission. Langmuir based models were also fitted to the binding dataset as described below. Error bars reported for all binding assays represent one standard deviation (±1σ) from mean values from three replicates.

***Langmuir-type adsorption model fitting to pull-down adsorption assay data:*** The bound (μmol protein/g cellulose) and free (μM) protein concentrations from the binding assays were fit to a Langmuir single-site, two-site, and Langmuir-Freundlich adsorption models to determine maximum binding capacity ($n_{max}$) and equilibrium dissociation constant ($K_d$) for each cellulose allomorph and GFP-CBM combination. The model equations are provided in **Figure 2C**. The linear range of the binding curve was used to estimate the partition coefficient ($\alpha$) as described previously (19). The data from pull-down binding assays is processed to yield free protein and bound protein concentrations (or calcofluor concentrations) in μM and μmol/g cellulose respectively in Excel™. This data set was then exported into OriginPro 2019. OriginPro software already has several non-linear curve fitting options available. However, separate functions were created for Langmuir single-site, two-site, and Langmuir-Freundlich models using the custom function builder tool. Similar Langmuir-type models have been used previously for Cel7A-cellulose binding analysis (2, 27). These functions were then used to fit binding assay data with the following settings. Levenberg-Marquardt algorithm with a tolerance of 1e$^{-9}$ was used for curve fitting. Fits did not converge in Origin for more complex models such as Langmuir two-site and Freundlich models. As a result, Excel Solver™ was used for curve fitting in those cases. The sum of errors was specified as objective function and parameters were estimated using the GRG non-linear method. Parameter uncertainties were then calculated using an elaborate process involving Monte Carlo simulations as detailed elsewhere (28).

***Reversible binding of GFP-CBMs to cellulose**:* The application of Langmuir-type adsorption models is dependent on reversible binding of protein to substrate. Hence, reversible binding properties of the GFP-CBMs to Cladophora cellulose I and cellulose III, as described previously (19, 29). Briefly, after the binding equilibrium was re-established after dilution of the mixture, the newly estimated bound/free protein concentration was confirmed to lie along the adsorption isotherm curve as already shown by us previously for CBM2a (ActE) (19). Since the partition coefficient was determined regardless of the dilution ratio, this result confirmed that the protein showed indeed reversible adsorption to cellulose. The presence of excess BSA prevented non-specific binding of CBMs to various surfaces like plastic tube walls (30), which along with reduced possible GFP-CBM denaturation at air-liquid interfaces during extensive mixing at low protein concentrations due to presence of sacrificial BSA (31), also minimized bias in reversibility binding measurements of GFP-CBMs particularly to cellulose III.

***Enzymatic hydrolysis of cellulose allomorphs*:** All Cladophora cellulose samples were subjected to enzymatic hydrolysis using a commercial cellulase cocktail at 0.5% glucan loading in a 1 ml reaction volume. All hydrolysis assays were carried out in 2 mL microcentrifuge tubes using 5 mg of pre-weighed cellulose suspended in 500 μL of 50 mM Na-Acetate buffer, at pH 5.0 (for C.Tec2), along with suitably diluted stock enzyme solution to achieve desired enzyme loadings (i.e., mg total enzyme loaded per gram of added cellulose per well). The total Cellic C.Tec2 enzymes (Novozymes, CA) loadings used during enzymatic hydrolysis was fixed at 5 mg/g glucan loading, unless specified otherwise. The protein concentration (193 mg/ml) for the C.Tec2 enzyme stock solutions was determined using the Kjeldahl method (32). Sodium azide was added to prevent any microbial growth (0.1% w/v final concentration). All tubes were incubated at 50 ˚C in an orbital shaking ThermoMixer (Eppendorf) incubator set at 1000 RPM for the desired saccharification time (0-96 h). The hydrolyzate supernatants were analyzed for total reducing sugar concentrations using the standard dinitrosalicylic acid (DNS) colorimetric assay as reported earlier (33). Briefly, 30 μl of the hydrolysate supernatant (w/wo 2-fold dilution) was incubated with 60 μl of DNS stock reagent in PCR tubes/plates at 95 °C for 5 min in an Eppendorf thermal cycler. After the PCR plates cooled down to room temperature, the DNS reaction mixture was transferred and diluted in DI water using a clear, flat-bottom microplate for finally measuring solution absorbance at 540 nm. Suitable reducing sugar standards (e.g., glucose standards ranging from 0.1–5 g/l) were included for the DNS assay. All hydrolysis experiments were carried out in duplicates. Error bars reported represent one standard deviation (±1σ) from mean values for replicate assays.

***Sample preparation for AFM imaging of Cladophora CI and CIII:*** Approximately 5 mg of dried cellulose I (CI) and cellulose III (CIII) fibers derived from *Cladophora glomerata* were each added to a microtube and suspended in 1 ml of DI water. At first, the fibers were manually dispersed through pipetting the suspension up and down using a wide opening 1 ml pipette. Subsequently, the suspensions were sonicated for 1 minute (model FB705 Fisher Scientific, USA, settings 10% amplitude, 2 seconds on, 5 seconds off), then pipetted up and down until a segregation of the fibers was observed. Two hundred microliters (200 µl) of the resulting suspensions were transferred to new microtubes and filled up to 1 ml with DI water. The fibers were further fragmented by pipetting through a 1 ml pipette until all large aggregates were dispersed. The suspensions were stored at 4°C until use and resuspended prior to usage.

***AFM imaging of Cladophora CI and CIII microfibrils:*** The microscope cover glasses (No. 1.5, 22x22mm, VWR, USA) were rinsed in the following order, DI water, acetone (NF/FCC grade, Fisher Scientific, USA) and DI water and then dried with a stream of nitrogen. Twenty microliters (20 µl) of cellulose I and III samples were each placed in the middle of the glass slide and dried over night at 50°C. Non-contact mode AFM measurements were carried out with a Park systems NX10 AFM using non-contact cantilever (SSS-NCHR, Park Systems, South Korea) that had a force constant of 42 N/m (specific range: 10-130) and a resonant frequency of 330 kHz (specific range: 204-497). For each of the substrates, 2.5 x 5 $\mu m^2$ sized areas were chosen at random places of the sample. Data were analyzed using the XEI software (Park Systems, South Korea).

***Preparation of Avicel cellulose nanocrystals through acid hydrolysis for QCM-D:*** Avicel cellulose III (or CIII) was prepared from Avicel PH-101 (also referred to as Avicel cellulose-I (or CI) (Sigma Aldrich Lot#BCBG9043V) as described previously (3). Nanocrystals were prepared from both Avicel cellulose-I and Avicel cellulose III using the same procedure. Briefly, 2 g of Avicel was added to 70 ml 4 N HCl in a glass beaker and placed in a pre-heated water bath at a temperature of 80$^o$ C. The slurry was stirred every half hour using a spatula to ensure cellulose is well suspended. After 4 hours of reaction time, the acid hydrolysis mixture was diluted with 50 ml deionized water. The slurry was then aliquoted into 50 ml centrifuge tubes, with 40 ml slurry in each tube and centrifuged at 1600xg for 10 minutes. The supernatant was decanted, and the cellulose pellet was washed with 10 ml deionized water. The wash steps were repeated, and the supernatants were discarded until they turned hazy around pH 3.3. The haziness of supernatant indicates evolution of cellulose nanocrystals and hence these supernatants were collected into a separate bottle for future usage.

***Preparation of cellulose thin films for QCM-D:*** This procedure was developed based on similar previous studies investigating cellulose hydrolysis and binding of cellulases using QCM-D (34). 4.95 MHz quartz crystal sensors (0.55" diameter), with $SiO_2$ coating were purchased from Filtech (product code QSX0303). The sensors were first rinsed in water, followed by ethanol and then blow-dried. The sensors were then immersed in 0.02% PDADDMAC (Sigma Aldrich 409022), which serves as an anchoring layer for cellulose, for 1 hour at 25$^o$C with orbital mixing. This was followed by washing with deionized water for 1 hour with orbital mixing at 25$^o$C. The sensors were then blow-dried and spin-coated using a pre-cycle spin for 3 seconds at 1500 rpm, followed by a spin cycle for 60 seconds at 3000 rpm. This spin coating step was repeated 10-20 times to obtain a uniform cellulose film thickness of ~20 nm (as measured using the QSoft software using Sauerbrey model). In a previous study (34), a lesser number of spin-coating steps was used to achieve the same thickness, however, this discrepancy may be related to the nanocrystal slurry concentration and hence needs to be optimized accordingly.

***Quartz crystal microbalance with dissipation (QCM-D) based CBM-cellulose binding assay and data analysis:*** Binding assays were performed using QSense E4 instrument (NanoScience Instruments). Quartz sensors with cellulose thin films were mounted and equilibrated with buffer (10 mM MES pH 6.5) at a flow rate of 100 µl/min for 10 minutes using a peristaltic pump. The flow of buffer was then stopped, and the cellulose films were left to swell overnight in buffer. The frequency and dissipation changes were tracked for all harmonics and the cellulose films were considered amenable to binding studies if the third harmonic stabilized after overnight equilibration in the buffer. Proteins of interest were diluted to a concentration of 1 µM beforehand and flown over the sensors at a flow rate of 100 µl/min for 10 minutes. All proteins tested, attained saturation within 10 minutes as noticed from frequency and dissipation traces. The CBM-cellulose system was left to equilibrate for at least 30 minutes. Unbinding of proteins was tracked by flowing 10 mM MES pH 6.5 buffer at 100 µl/min for at least 30 minutes. The sensors were finally treated with 5% contrad solution followed by deionized water at 100 µl/min for 10 minutes, to remove any traces of protein left in the tubing. The frequency data was converted to mass deposited on sensor using Sauerbrey equation, which was in turn converted to the number of molecules of protein deposited on cellulose surface. This data was then used to obtain pseudo-association rate constant ($K_{on}$*) and dissociation rate constant ($K_{off}$) using the equations provided below:

<u>Binding rate equation:</u>

$$[EC] = A(1 - e^{-(K_{on}^{*})t})$$

<u>Unbinding rate equation:</u>

$$[EC] = Ae^{-(K_{off})t}$$

Where $[EC]$ = Number of molecules; $t$ = Time (minutes)

***Fluorescence recovery after photobleaching (FRAP) to study CBM-cellulose binding kinetics:*** We assembled a simple fluidic chamber using a glass coverslip (24 mm X 30 mm, No. 1.5, Thermo Scientific) and a holed glass slide (470150-480, Ward's Science) that was pre-cleaned with acetone (A949, Fisher chemical), methanol (A454, Fisher chemical), and distilled water. To improve the cellulose sample adhesion, the glass substrates were treated with plasma (PDC-001, Harrick Plasma) for 10 minutes. A drop of 20 μL cellulose solution was deposited on a coverslip, which was dried in a hybridization oven (VWR) at 50°C overnight. A sample chamber was assembled by attaching the coverslip to a glass slide with multiple pieces of double-sided 3M tape; the gap between the edges of two adjacent tapes formed a channel. The open edges were sealed with 5 minutes epoxy glue (Devcon). Next, 100 μL of 5 μM GFP-CBM3a buffered solution (10 mM MES pH 5.5) was finally injected into the sample chamber through the holes on the glass slide, which then was incubated for 10 minutes before running FRAP experiment.

For FRAP experiment, we used a Total Internal Reflection Fluorescence (TIRF) microscope, which was homebuilt with an inverted microscope (Ti-E, Nikon), high NA objective lens (CFI-apo 100X, NA 1.49, Nikon), 488 nm laser (Coherent) and EMCCD camera (iXon Ultra-888, Andor) (35). We scanned through the cellulose sample to find an area to be imaged and acquired a reference wide-field fluorescence image at a low 488 nm excitation power (2-3 mW) prior to photobleaching. Guided by the pre-bleach image, we decided on multiple locations to be photobleached, and created focused laser spots at the corresponding positions using a spatial light modulator (X13138-01, Hamamatsu). Typically, a maximum of 25 spots on the cellulose fibers along with 2 spots on the background area were chosen. We locally photobleached the selected spots with illumination of a strong 488 nm laser power (50-120 mW) for 1 minute, and then time-lapse wide-field fluorescence images were acquired for 40 minutes to monitor fluorescence recovery. Focus drift was actively compensated with Perfect Focus System (Nikon) while XY-drift was corrected using ImageJ software (NIH).

Custom written MATLAB scripts were employed for the extraction of the recovery curves and data analysis. The photobleached segments were automatically identified by subtracting pre-bleach from post-bleach images. The recovery curves were corrected for non-specific GFP-CBM binding to the glass slide within a segment by setting all pixel values below a segment-specific threshold to zero. The average pixel intensity value of each segment before photobleaching was used to normalize the recovery curve. Additionally, the FRAP signal was baselined to zero relative intensity based on the first FRAP data point to account for the non-zero dark current of the camera and inhomogeneous illumination across the FOV as well as inefficient photobleaching. Each FRAP curve was fitted to the model given in equation below developed by Moran-Mirabal (36) using a Levenberg–Marquardt curve fitting approach built-in MATLAB. The baseline correction is then added to $F_{M,fit}$ to obtain the true $F_M$ value.

$$I(t) = F_{M,fit} * \left(1 - e^{k_{off} * t}\right)$$

With $I(t)$ being the normalized intensity value at time $t$, $F_M$ being the fraction of reversible bound GFP-CBM3a, and $k_{off}$ the unbinding rate of GFP-CBM3a assuming a first order binding reaction. Fits with an $R^2 \leq 0.85$, $F_M \leq 0$ or $F_M \geq 1$ were excluded from further analysis. The collection of fit parameters for each protein and cellulose allomorph combination $(F_M, k_{off})$ were then fitted to a Gaussian distribution to extract the mean and standard deviation of each parameter.

***Functionalization of beads for tweezer binding-rupture and motility assays:*** CBM1 was tethered to polystyrene beads via the $His_8$-tag on the N-terminus of our purified GFP-CBM1 construct, with minor modifications from our previously published work (1). Using PCR, 1,010-bp DNA linkers were created from the M13mp18 plasmid template with a biotin tag on one end and an amine group on the other. The anti-His antibody was crosslinked to the amine group using a sulfo-SMCC intermediate. In the cases of using the anti-His Fab, the anti-His antibody was cleaved using 3-MEA before crosslinking. To functionalize the beads with GFP-CBM1, 1.09 μm streptavidin beads (Spherotech), biotin/anti-His functionalized DNA linkers, and $His_8$-tagged GFP-CBM1 constructs were incubated together in PBS at 4°C for 45 minutes on a rotator. After incubation, the beads were washed by spinning down at 9000 rpm for 3.5 minutes, removing the unreacted components in the supernatant, resuspending in 50 mM acetate buffer (pH 5.0), and sonicating for 2 minutes at 20% amplitude. This process was repeated two more times. Beads were functionalized such that, statistically, zero or one GFP-CBM1 molecule is bound to each bead. This was determined through serial dilution until a maximum of half the beads bound to cellulose fibers during the experiment. Lastly, purified Cel7A enzymes were tethered to polystyrene beads and assayed for their motility on Cladophora derived cellulose I and cellulose III based on identical methods reported earlier in our Cel7A tweezer motility study (1).

***Cellulose solution and slide preparation for tweezer binding-rupture and motility assays:*** Purified and dried cellulose samples (*Cladophora* based cellulose I or III) were used to create a heterogeneous cellulose mixture by first mixing the desired cellulose sample to deionized water in a 1 mg/mL ratio. The mixture was then sonicated for 2 minutes at 50% in a cup sonicator and vortexed for 15 seconds on high setting. The cellulose, still clumped at this point, was pulled up and down in solution with a 16-gauge syringe for 1-2 minutes before going back on the vortex for 15 seconds. These steps were repeated three times. The resulting mixture was then diluted in a 1:20 ratio by mixing 500 µL of the prepared solution with 500 µL deionized water. This slurry suspension was then stored at 4 °C. Whatman Grade 1 Filter Paper based cellulose stock suspension slurry was prepared as described previously (1), to be used for some control GFP-CBM1 binding-rupture assays. When preparing to load a slide, a small sample (~100 µL) of the stored cellulose mixture is removed from the stock and the cellulose pulled apart by sonicating for 2 minutes at 50% in a cup sonicated. This solution was directly loaded onto the glass slide. Slides are prepared by creating a 10-15 µL volume flowcell using a KOH etched coverslip and double-sided sticky tape. The stock cellulose solution (*Cladophora* based cellulose I or III) was then added to the flowcell and allowed to dry out in an oven at ~95 °C for an hour, allowing cellulose fibrils to non-specifically bind to the slide surface. The surface was then blocked with 10 mg/mL BSA in acetate buffer (pH 5.0) for 15 minutes to prevent non-specific sticking of the beads to the glass surface. Finally, the GFP-CBM1 functionalized beads solution was loaded onto the slide and the slide sealed shut. For the Cel7A motility assays, 0.75 $\mu$m non-functionalized polystyrene beads (Spherotech—PP-08-10) were allowed to nonspecifically adhere to the coverslip surface, in an incubation step before BSA blocking, to serve as fiducial markers allowing for instrumental drift tracking during data acquisition.

***Single molecule tweezer binding-rupture assay data acquisition and analysis:*** CBM1 functionalized beads were trapped using a 1064-nm laser setup as described before (1), and placed alongside a surface-bound stationary fiber. Experiments were conducted at a fixed room temperature (21 °C). After position calibration and trap stiffness measurements, the bead was actively placed on a cellulose fiber roughly running along the axis of the microscope stage. Upon binding, the bead was centered, acquisition started, and a force applied to the tethered bead by stepping the piezo stage along the axis of the fiber. With force applied, the position of the bead is held until rupture. Once a tether is ruptured, it is sometimes possible to tether the bead to the fiber again, in which case, the same method of force application is applied while data acquisition continues. Data were collected at a 3-kHz sampling frequency and then filtered with a 10-point exponential moving average before analysis. Custom Matlab codes were then used determine the rupture forces and the bond lifetimes of full ruptures. The force-lifetime data was binned every 2.5 pN and then we tried to fit the data to a single or a double exponential decay characteristic of a slip bond.

***Molecular dynamics simulation of CBM1 and cellulose allomorphs interactions:*** All molecular dynamics (MD) simulations were conducted using the Amber force field for CBM1 protein (1cbh pdb code) (37), TIP3P explicit water model (38), and the Glycam force field for representing the cellulose fiber (39). As described previously (40), X-ray and neutron diffraction coordinates were used to generate one rhomboid cellulose Iβ fibril and one rhomboid cellulose III fibril (11, 12). Each fibril consisted of 36 glucan chains each. Two different types of crystalline cellulose III were used for the simulations, one with lower crystallinity and another with high crystallinity (based on the available crystalline core cellulose crystal structure). However, since no detailed structural information is available for lower crystallinity cellulose III (3), an idealized model structure was obtained by increasing the simulation temperature to 400 K starting with the high crystallinity cellulose III structure to increase overall disorder within the crystalline fiber. The cellulose Iβ rhomboid shape was chosen as the base case control because of its wide hydrophobic surfaces accessibility which has shown to be preferred binding site for Type-A CBMs like CBM1 (41, 42). Furthermore, this hydrophobic binding surface plane of cellulose Iβ is also identical to cellulose I$\alpha$ (10), from a crystallographic point of view, which would aid in drawing similar conclusions when assessing CBM1 binding to hydrophobic surfaces of native cellulose I fibrils from Cladophora, which is mostly enriched in cellulose I$\alpha$ unlike cellulose I fibrils from Avicel derived cellulose nanocrystals that are enriched in cellulose I$\beta$.

Two types of simulations were run: (a) First, unbiased simulations were used to probe the binding dynamics of CBM1 starting from the hydrophobic surfaces of cellulose I and III microfibrils on microsecond timescales. MD simulations were conducted in order to populate the most preferred orientations of CBM1 on either cellulose allomorph surface. CBM1 was first aligned in the canonical direction for the reducing end specific action of Cel7A to processively hydrolyze cellulose chain (i.e., the Y31 residue end of CBM1 pointing towards the nonreducing end of cellulose). If no stable binding was seen, as was the case with cellulose III, the CBM1 was aligned in the most stable conformation identified in the unbiased simulations (i.e., Y31 residue end of CBM1 pointing towards the reducing end of cellulose). From these unbiased simulations, we were able to calculate the distribution frequency of the most preferred rotameric states for the closely cellulose-contacting aromatic residues in CBM1. (b) Second, the preferred orientation inferred from unbiased simulations for CBM1 on cellulose I and III allomorphs were used in order to estimate their respective association free energy with the cellulose fiber. Thus, a potential of mean force (PMF) was calculated to estimate the CBM1 binding free energy during adsorption to the hydrophobic surface of all cellulose allomorphs. We used the coordinates of the last frame from the unbiased CBM1-cellulose systems at 310 K. A total of 14 independent windows per system were used, which were spaced apart by 1 Å. A restraining potential of 1000 kJ $mol^{-1}$ $nm^{-2}$ was applied to the center of mass (COM) of CBM1 with respect of the COM of either cellulose I or cellulose III along the normal (z) coordinate. For each window, 1.5 µs long simulations were performed. The desorption free energy was reconstructed using the weighted histogram approach and convergence was assessed via block averaging by dividing the trajectory in three independent blocks. Umbrella sampling-based simulations were conducted along a path where the CBM1 translated from the hydrophobic surface of the microfibril to the bulk solvent. The position of the potential changes every 1 Å along the pulling reaction coordinate (in this case the z vector). Harmonic potential applied to the center of mass of the protein. Along the trajectory, the two most populated configurations are shown here in the main text figure, together with the protein residues in close contact with the external cellulose surface. These configurations also correspond to the minimum wells seen in the energy plots.

All simulations were performed using a 2 fs time step using GROMACS software. The LINCS algorithm was applied to constrain all bond lengths with a relative geometric tolerance of $10^{-4}$. Non-bonded interactions were handled using a twin-range cutoff scheme. Within a short-range cutoff of 0.9 nm, the interactions were evaluated every time step based on a pair list recalculated every five-time steps. The intermediate-range interactions up to a long-range cutoff radius of 1.4 nm were evaluated simultaneously with each pair list update and were assumed constant in between. A PME method was used to account for electrostatic interactions with a grid spacing set to 0.15 nm. During the equilibration (0.1 us), systems were coupled using a Berendsen barostat to 1.0 bar via an isotropic pressure approach, with relaxation time of 1.0 ps. Afterwards, system was coupled to a Parrinello barostat algorithm and constant temperature was maintained by weak coupling of the solvent and solute separately to a velocity-rescaling scheme with a relaxation time of 1.0 ps.

***Buffon needle model analysis of CBM orientation probability distribution on cellulose surface:*** Buffon's needle model arose from a problem first posed by the eponymous French mathematician Buffon (43, 44). The model is an analytical solution to the question of the probability of a short needle crossing the parallel lines on a ruled paper or wooden floor. An analytical problem to this solution does exist and the numerical solution to this problem would involve the Monte Carlo method. Here, we have adopted the Monte Carlo approach to simulate the CBM-cellulose system with some simplifications which render the binding process completely geometric without any energetic constraints. The CBM was assumed to be a needle with length of 20.7 Å since the distance between centers of flanking planar Tyrosine rings (Y5 and Y31) is 20.7 Å based on the published NMR structure (1cbh PDB). Cellulose-I was assumed to be an array of lines, each line representing a cellulose chain, with an inter-chain distance of 8.2 Å. In the energetically favorable configuration of this system, CBM1 aligns with two aromatic residues (Y31, Y32) on one cellulose chain and another aromatic residue (Y5) on an adjoining cellulose chain (see **Fig. S16**). Hence, any event where the CBM needle lands on a chain perfectly ($C_0$), crosses only one chain ($C_1$) were clustered into a category where the CBM can be interacting only with two cellulose chains. Similarly, events where the CBM needle crosses two chains ($C_2$) and three chains ($C_3$) were clustered into a category where the CBM can be interacting with more than two chains. The configuration of this system at any time can be defined by the angle ϴ between the cellulose chains and the CBM needle. A Monte Carlo simulation was then conducted with 10000 trials whereby a random number generator was used to sample the angle ϴ. Configuration from a given trial can then be classified into one of the five categories mentioned above ($C_0$ – $C_3$) and eventually, the probability of CBM crossing two chains or more. See **Fig. S16** for additional methodology related details.

**SI Appendix Results and Discussion**

***Cellulose allomorph characterization by XRD and Raman Spectroscopy**:* Cellulose crystallinity index based on the Segal method was estimated to be about 90-95% for both allomorphs (SI Appendix **Fig. S1A**). Cellulose crystallite size was about 8.5-9 nm for both allomorphs, estimated using the Scherrer equation based on the full-width half-maximum of the 200 equatorial plane reflection peaks. Cladophora cellulose based crystallites were at least 2-3 times larger in cross-sectional diameter than previously reported for cellulose microfibrils derived from higher-order plants, such as cotton linters (2). Similar to previous reports (2, 16, 17), Raman spectroscopy also independently confirmed that native Cladophora cellulose I was completely converted into cellulose III following ammonia treatment (SI Appendix **Fig. S1B**).

***AFM imaging of Cladophora cellulose allomorphs:*** Previous studies on cellulase or CBM binding have often employed two-site binding models (45), with the assumption of two distinct classes of binding sites arising from any of the following phenomena: (i) binding to amorphous/crystalline regions or (ii) binding to two different faces of crystalline cellulose fiber or (iii) due to different orientations of CBM on hydrophobic face of cellulose. Since Cladophora derived cellulose is a highly crystalline substrate and Type-A CBMs bind to crystalline regions predominantly, a small number of high affinity amorphous binding sites seems unlikely. However, the availability of different crystal faces for binding to non-native allomorph cellulose III has not been examined before and hence we performed AFM imaging on both Cladophora cellulose I and cellulose III microfibrils to verify the general morphological differences between two substrates (see **Fig. S6**). AFM imaging indeed suggests that the Cladophora cellulose I elementary fibrils have two distinct crystalline planar surfaces likely available for CBM binding unlike cellulose III, consistent with predicted crystal structure of algal cellulose and morphological changes undertaken following ammonia treatment (10, 12, 46). This change in cellulose I fibril shape is expected to take place during ammonia pretreatment due to a solid-state polymorphic transformation that alters the cellulose III unit cell crystal structure (12, 47–49). Hence, heterogeneity in binding sites for cellulose I versus cellulose III could arise from other reasons and any physical interpretation of CBM-cellulose binding interactions based solely on model fit parameters could be misleading.

***Type-A CBM3a exhibits increased desorption rate constant towards cellulose III:*** Classical solid-state depletion assays, as discussed above, confirmed that a 2-13 fold lower CBM3a binding affinity is observed towards Cladophora derived cellulose III depending on whether a one- or two-site Langmuir adsorption model is applied for data analysis. However, these assays do not provide any information on the relative impact of the association versus dissociation rate constants on the equilibrium binding affinity constant. Estimation of association and dissociation rate constants could further yield insights into the rate-limiting steps for CBM adsorption to cellulose allomorphs and cellulase-mediated degradation of cellulose in general. We developed and applied two different methods to study CBM-cellulose binding kinetics, one employing fluorescence recovery after photobleaching (FRAP) (36, 50) and another employing quartz crystal microbalance with dissipation (QCM-D) (51, 52). CBM3a was picked for this study, since it exhibited a clear 3-fold reduction in bulk binding affinity towards cellulose III for the one-site model (see **Table 1**) and hence potentially allows for a significant difference to be observed in the association and/or dissociation rate constants. In addition, we used FRAP to study GFP-CBM3a binding to Cladophora derived cellulose allomorphs (with native cellulose enriched in I$\alpha$) and QCM-D to study GFP-CBM3a binding to plant-derived Avicel microcrystalline cellulose allomorphs (with native cellulose enriched in I$\beta$) (53). Hence, this would present a more comprehensive picture of a model Type-A CBM binding kinetics to cellulose III allomorphs obtained from two distinct biological sources.

Procedures for preparation of cellulose nanocrystals, nanocellulose thin film deposition, QCM-D binding assay, and data analysis procedures are outlined in the SI appendix materials and methods section. Sauerbrey equation was used to obtain the mass of adsorbed protein on cellulose film using the frequency change at third overtone (54). The number of protein molecules was then estimated from the mass of adsorbed protein and this resulted in QCM sensorgrams as shown in **Fig. S8A-B.** This data was then analyzed using exponential fitting routines (see exponential fits to acquired QCM raw data in **Fig. S8C-D**), to obtain a pseudo association rate constant ($K_{on}$*) and dissociation rate constant ($K_{off}$). While $K_{on}$* did not show a significant difference between the two allomorphs, CBM3a gave a nearly 3-fold increase in $K_{off}$ for cellulose III (**Table S1**). In addition, the maximum amount of protein bound to cellulose III was 1.5-fold less.

These observations align well with the reduction in binding of CBM3a to cellulose III observed using the classical pull-down binding assay method. In addition, frequency (third overtone) was plotted against dissipation (third overtone) as shown in **Fig. S8E.** The protein layer adsorbed on cellulose III showed up to 20-fold higher dissipation compared to cellulose I. This indicates the formation of a 'softer' or less viscoelastic adsorbed protein layer on cellulose III, likely arising from the increased diffusion of bound proteins. Similar QCM-D results during CBM binding to another allomorph (i.e., cellulose-II) have been reported as well (55).

Next, we performed FRAP experiments, with details regarding the assay method and data analysis outlined in the SI appendix materials and methods section. Briefly, similar to previous work (36, 45), GFP-CBM3a binding kinetic parameters to cellulose allomorphs were obtained by fitting the FRAP curves to a binding-dominated model ignoring any diffusion relevant contributions. Our FRAP analysis revealed that CBM3a gave a 1.9-fold increase in $K_{off}$ for cellulose III compared to cellulose I (**Fig. S9**). In summary, solid state depletion equilibrium binding assays as well as both QCM and FRAP binding assays together provide complementary information on the differences in binding affinity and kinetic rate constants for CBM binding to cellulose allomorphs. However, neither of these methods alone provide a detailed molecular-level understanding of Type-A CBM binding, particularly for weaker binders like CBM1 that impact cellulase binding and activity towards distinct allomorphs.

***Interpretation of equilibrium binding parameters and sensitivity analysis:*** We varied the free protein concentrations over nearly four orders of magnitude (0.02-250 μM for CBM1 and 0.02-50 μM for CBM3a) to accurately capture binding interactions at both low (<10-fold of min $K_d$) and high (>10-fold of max $K_d$) protein-to-substrate saturation concentrations. This broad range of protein concentrations is often recommended to properly characterize CBM/CAZyme adsorption to polysaccharides (56), but is not always followed by researchers, which could explain some of the discrepancy in the literature for reported $n_{max}/K_d$ values for various CBMs. Langmuir two-site models are often interpreted as consisting of two different classes of binding sites (45), one with high affinity and another with low affinity (a higher value of $K_d$ corresponds to a lower affinity). Langmuir-Freundlich model, on the other hand, also provides information regarding the binding cooperativity for a given substrate (57). However, since most studies do not validate such model assumptions, it is very challenging to draw any molecular-level understanding of protein adsorption process using such traditional approaches alone. Furthermore, since the root-mean square errors from the model fits to the experimental data are nearly comparable in most cases, it is difficult to choose an appropriate model based on goodness of model fit without any bias. Details regarding sensitivity of results to truncation of original CBM1 data set is discussed already (see **Table S2**). Furthermore, we also performed sensitivity analysis for the original CBM1 to cellulose I binding data set by studying the variation in RMSE when the binding parameters ($n_{max}, K_d$) are varied individually by 10% and 20% each (see **Table S3**). This sensitivity analysis revealed that the $n_{max}$ parameter is quite sensitive to error in data acquired at higher protein concentrations, emphasizing the need for enough replicates while probing higher protein concentrations.

***MD simulations reveal improper stacking interactions of aromatic residues with cellulose III:*** MD simulations were conducted to identify the preferred conformations of CBM1 on each cellulose allomorph surface and estimate the theoretical binding affinity of CBM1 for the most preferred orientation. Two different types of crystalline cellulose III were used for the simulations, one with lower crystallinity and another with high crystallinity (based on the available crystalline core cellulose crystal structure). A recent study from our group (3) has shown that CBM3a binding to cellulose-III closely depends on the fibril crystallinity. Therefore, we were interested in characterizing the binding of CBM1 to two types of cellulose III crystals with varying degree of crystallinity for this study as well. However, since no structural information is available for lower crystallinity cellulose III, a modeled structure was obtained by increasing the simulation temperature to 400 K starting with the high crystallinity cellulose III structure to increase overall disorder within the crystalline fiber. Previous studies employing MD simulations (42) and CBM-cellulose EM-immunolabeling experiments (58) have already shown that the preferred face for binding of CBM1 to cellulose I is the hydrophobic face. Therefore, the hydrophobic faces of cellulose I and cellulose III were initially chosen to study the binding preferences of CBM1 in this study as well.

As shown in **Fig. S14A**, the unbiased MD simulations revealed that the most populated configurations for CBM1 on cellulose I have the CBM1 Y31 residue facing towards the non-reducing end of the cellulose chain (as also reported previously (41)). This chain-end preference could have evolved from the reducing end specificity of the Cel7A catalytic domain to which the native CBM1 is attached at the C-terminus. Furthermore, it was also observed that CBM1 remained stably bound to cellulose I surface regardless of whether the Y31 residue was facing the reducing or non-reducing end. Interestingly, CBM1 was stably bound to the cellulose III surface (for both low and high crystallinity forms of cellulose III) only when the Y31 residue was facing the reducing end of cellulose. See **Movies S1-S3** for representative visualization of CBM1-cellulose allomorphs interactions from unbiased MD simulations. The relative diffusivity of CBM1 and the RMSF for key aromatic residues is provided here as well (see **Fig. S14B-C**).

***Reduced calcofluor dye adsorption to cellulose allomorphs**:* Calcofluor is a trans-stilbene based derivative fluorescent dye that has a low molecular weight (917 g/mol), but has a core structure analogous to the planar interface of Type-A CBMs involving both aromatic and polar functional groups that provide the thermodynamic driving force for binding to multivalent cellulose surfaces. Calcofluor is thus an ideal inorganic CBM-like surrogate probe useful for understanding the multivalent binding interactions. Interestingly, Calcofluor dye was also found to show reduced partition coefficient towards microcrystalline cellulose III by 2.6-fold versus native cellulose I (see **Fig. S15A**). Calcofluor is expected to bind cellulose with high affinity, based on reports on how it impacts *in vivo* cellulose synthesis (59) and histological staining analysis of cell wall polysaccharides (60). However, to the best of our knowledge, a direct measurement of calcofluor affinity to microcrystalline cellulose has not been reported. Based on our two site Langmuir model fitting analysis, we found that calcofluor white dye binds to microcrystalline cellulose I with a lower $K_{d1}$ of 38.8 μM to 81% of total available sites and $K_{d2}$ of 1.2 μM to the remaining 19% of total available sites ($n_{max}$ total was 21.5 μmol/g cellulose). For cellulose III, we found that calcofluor white dye binds with a higher $K_{d1}$ of 79.3 μM to 70% of total available sites and $K_{d2}$ of 1.8 μM to the remaining 30% of total available sites ($n_{max}$ total was 9.2 μmol/g cellulose). Based on single site model fitting analysis, we found that calcofluor white dye binds to microcrystalline cellulose I with a $K_d$ of 18±3 μM and $n_{max}$ of 19.9±1.0 μmol/g cellulose, while for cellulose III a $K_d$ of 15±4.5 μM and $n_{max}$ of 7.7±0.5 μmol/g was estimated. However, the two-site model gave a much better fit for the data unlike the one-site model. Three-site model gave no further improvement to model fitness for either cellulose I or III. While, these values are likely dependent on the buffer ionic strength conditions, calcofluor clearly has lower affinity and binding sites available for adsorption to microcrystalline cellulose III versus native cellulose I. It is known that strong non-covalent interaction forces stabilize interaction of calcofluor along the repeating ~1 nm cellobiosyl-unit of cellulose along the chain axis (60). This could explain why calcofluor dye binds in a well-defined orientation parallel along the fiber axis to both chitin and cellulose (61). However, altering the crystal structure of native cellulose I to cellulose III could destabilize calcofluor binding due to steric clashes with adjacent cellulose chains due to changed crystal morphology or increased fibril hydrophilicity. These findings shed further light into the thermodynamic mechanism driving reduced binding interactions of critical planar CBM binding surface associated amino acid residues to cellulose III. Interestingly, as reported previously for CBM1 binding to crystalline cellulose (57), Scatchard plot analysis for calcofluor binding to cellulose was also non-linear and concave-upward (see SI appendix **Fig. S15B**). Calcofluor dye binding to multiple classes of non-equivalent binding sites could provide a classical interpretation of concave-upward Scatchard plots (62). Nevertheless, when characterizing bulk ensemble binding interactions to highly heterogenous crystalline cellulose microfibril surfaces, even for simple CBM-analogues like calcofluor, our work brings to the light the severe challenges associated with choice of multi-site adsorption models and possible misinterpretation of results.

***Buffon needle model for predicting CBM1 orientations:*** Here we applied a simple 'Buffon needle-type' geometrical probability type model (ignoring any energetic barriers) to gain insights into the theoretical orientations of CBM1 'needle' along or across multiple glucan chains on the cellulose surface. Buffon's model calculates the theoretical probability distribution P(x) for a needle (of length 'L') to lie across a line between parallel line strips (each line separated by width 'd') or at distinct angles (θ). Here, in our version of the model we have approximated the CBM1 planar aromatic binding residues motif length to be equivalent to the total Buffon needle length, the distance between individual cellulose polymer chains to be equivalent to the parallel lines, and the angle of the fallen needle with respect to the parallel lines to be equivalent to the CBM orientation along or across the cellulose chains. The objective was to estimate the frequency of events whereby the CBM would align along one cellulose chain as opposed to aligning across multiple cellulose chains. CBM1 is represented as a needle of 20.7 Å length (based on the distance between terminal aromatic Y5 and Y31 residues – see **Fig. S16**) whereas the cellulose surface is simply represented as a grid of parallel lines with a spacing of 8.2 Å between individual cellulose chains. A customized MATLAB code was then used to randomly sample the angle ($\theta$), and for each orientation specified by the angle $\theta$ the total number of cellulose chains crossed by the CBM1 needle were quantified. The Buffon model predicts that the probability of a CBM1 'needle' to bind along a single cellulose chain is ~40%, while the remaining ~60% of events would include binding across multiple cellulose chains (ignoring energetic barriers). Interestingly, if a mutation (Y31A for instance) is considered as having reduced needle length, that would increase the percentage of events along the chain to 90%. Hence, performing these planar aromatic residue mutations and testing the impact of these mutations on the heterogeneity of CBM binding to cellulose studied using bond rupture assays could give us some insight into the role played by CBM binding orientations on observed heterogeneity. Preliminary data from our CBM-cellulose bond rupture assay indeed shows that there are some distinct differences in the force-lifetime plots for CBM1 wild-type and Y31A mutant on cellulose **I** (e.g., see 0-2.5 pN and 10-15 pN ranges in F**ig. S13**). However, it needs to be emphasized that this is an over-simplistic model and an understanding of the energetic constraints could better simulate the complex reality of CBM-cellulose interactions. Most previous CBM binding focused studies (41, 42, 63) have not emphasized the possible orientations of CBM1 on the surface of cellulose **I**. Beckham *et al.* (41) previously showed that although CBM1 prefers to bind along the cellulose chain as well, slightly rotated (by ~10-15°) CBM1 orientations across multiple cellulose chains are energetically feasible as well on individual fiber surfaces. More detailed MD simulations and corresponding rupture assays need to be conducted to check how mutations of CBMs impact along the chain axis versus across multiple chain axis binding and its potential impact on non-productive processive cellulase binding.

**SI Appendix Figures & Tables**

**Fig. S1.** XRD (A) and FT-Raman (B) spectra for Cladophora-derived Cellulose I and Cellulose III.

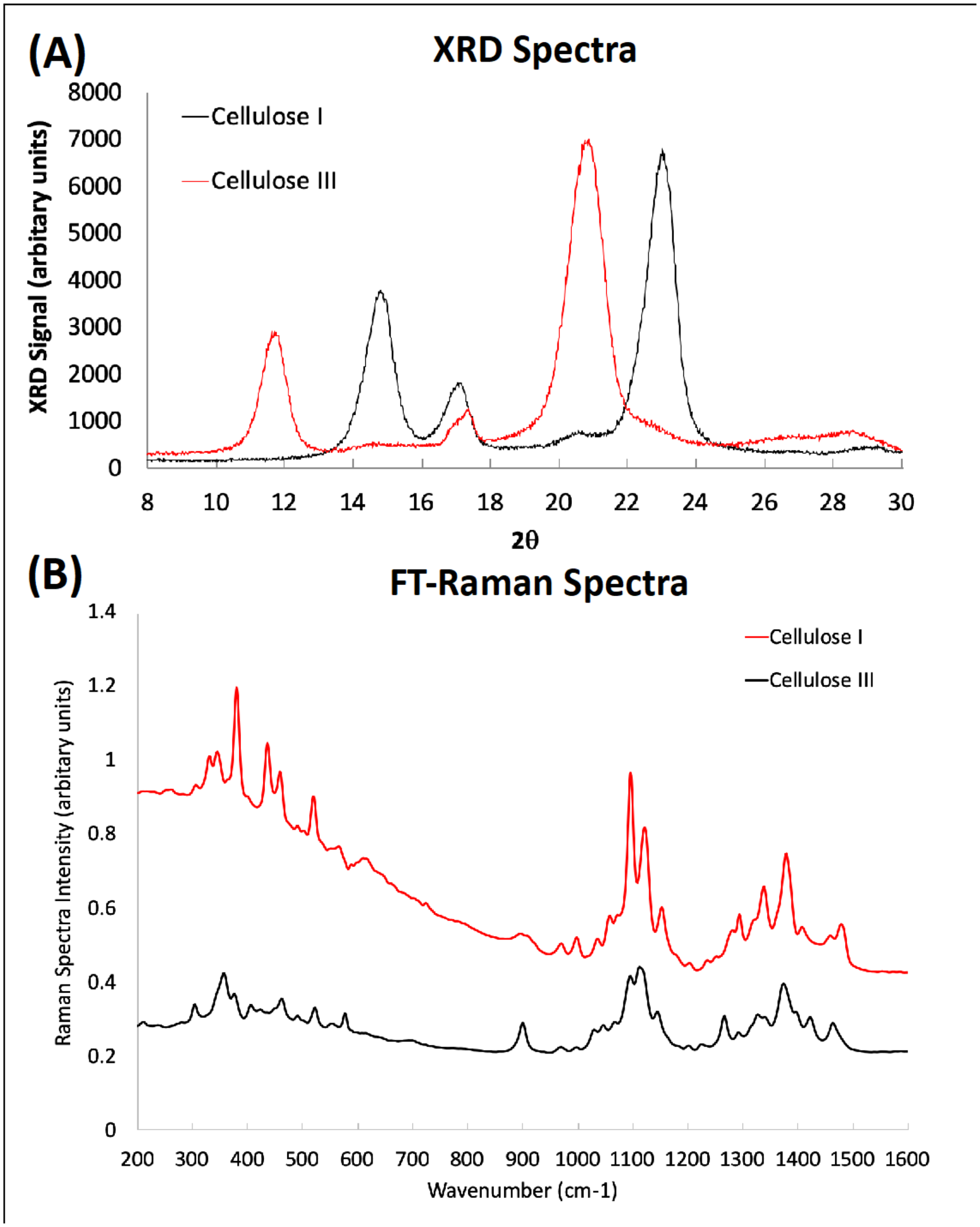

**Fig. S2.** Commercial fungal cellulase cocktail (C.Tec2) catalyzed hydrolysis of Cladophora-derived Cellulose I and Cellulose III for two different saccharification time points (24 and 96 h). Error bars here depict one standard deviation from mean for replicate assays.

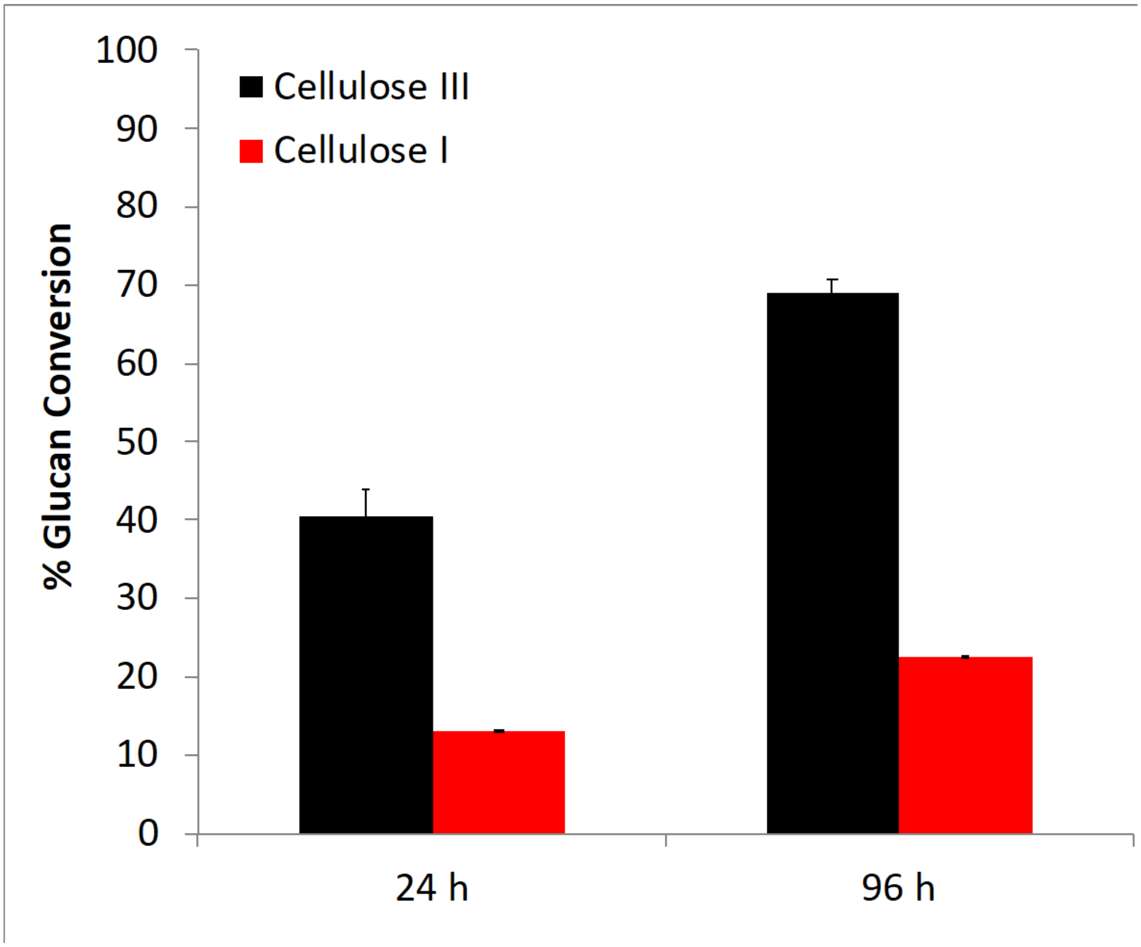

**Fig. S3.** Single-molecule optical tweezers-based verification of binding stability and instability for full length Cel7A cellulases processively deconstructing Cladophora based cellulose I and III. Three representative traces each are shown for the conditions of stability on cellulose I (red), stability on cellulose III (blue) and instability on cellulose III (black). Cel7A was found to be more often stably bound to cellulose I than to cellulose III before initiating stable processive motility accordingly. Of the traces showing instability, Cel7A was likely to have more rupture events on cellulose III than on cellulose I.

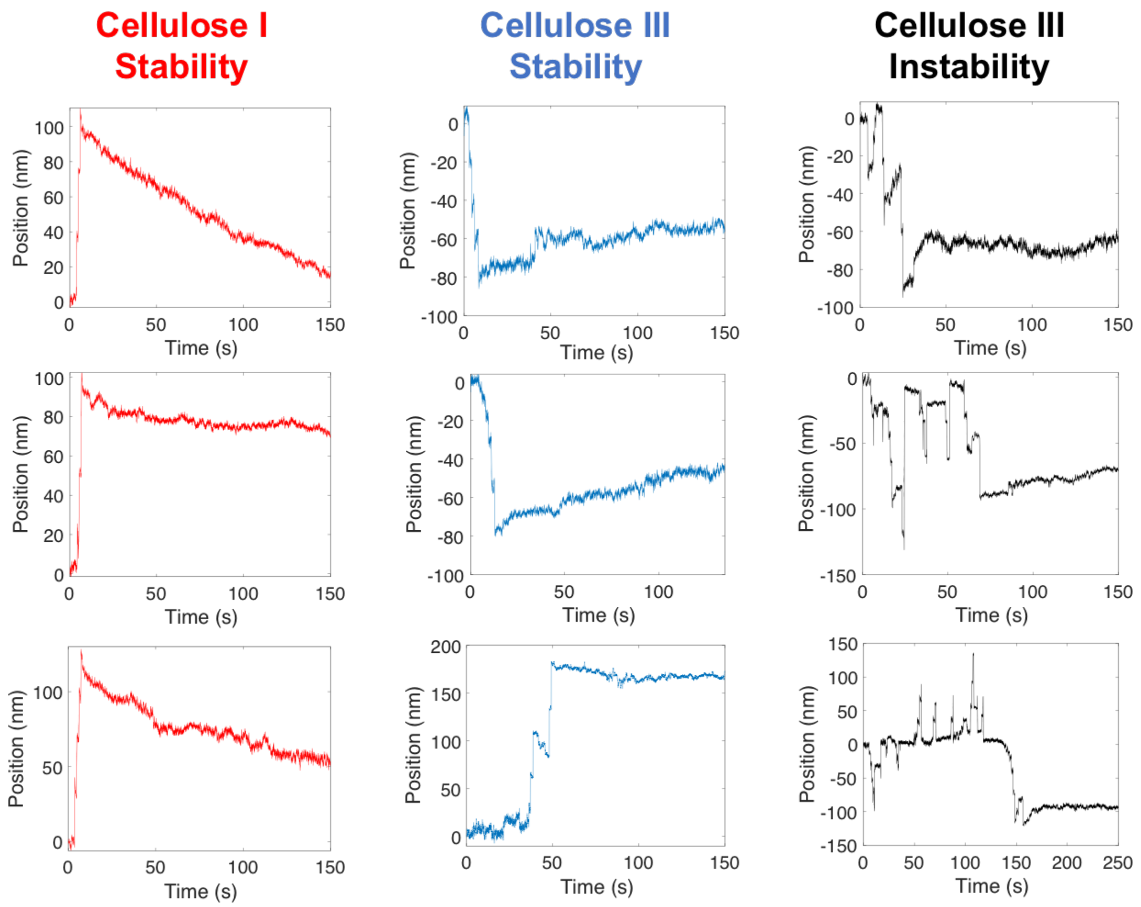

**Fig. S4.** Langmuir-type adsorption model fits (in red) for CBM1 binding data (in black) to Cladophora-derived Cellulose I (A, C, E) and Cellulose III (B, D, F).

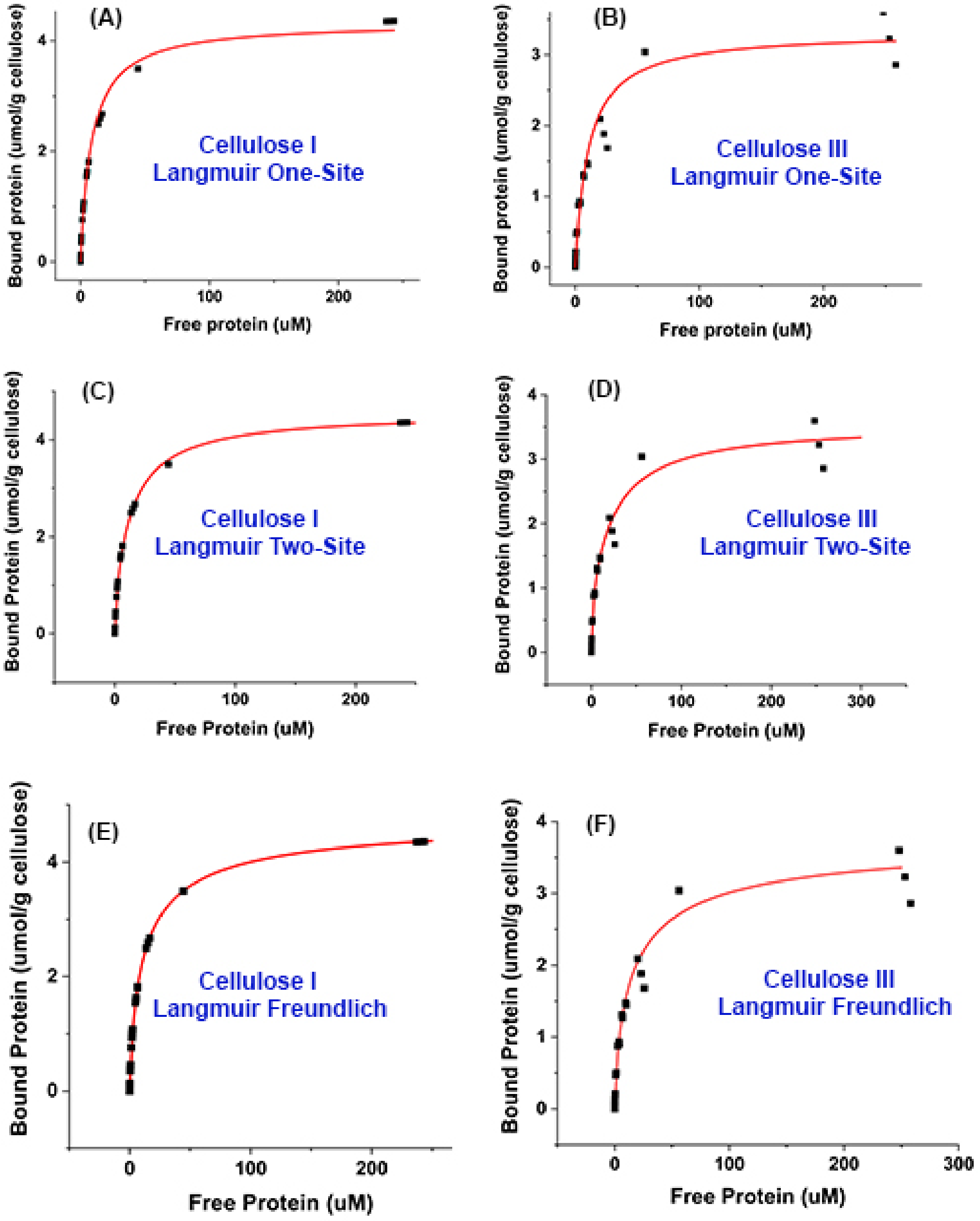

**Fig. S5.** Langmuir-type adsorption model fits (in red) for CBM3a binding data (in black) to Cladophora-derived Cellulose I (A, C, E) and Cellulose III (B, D, F).

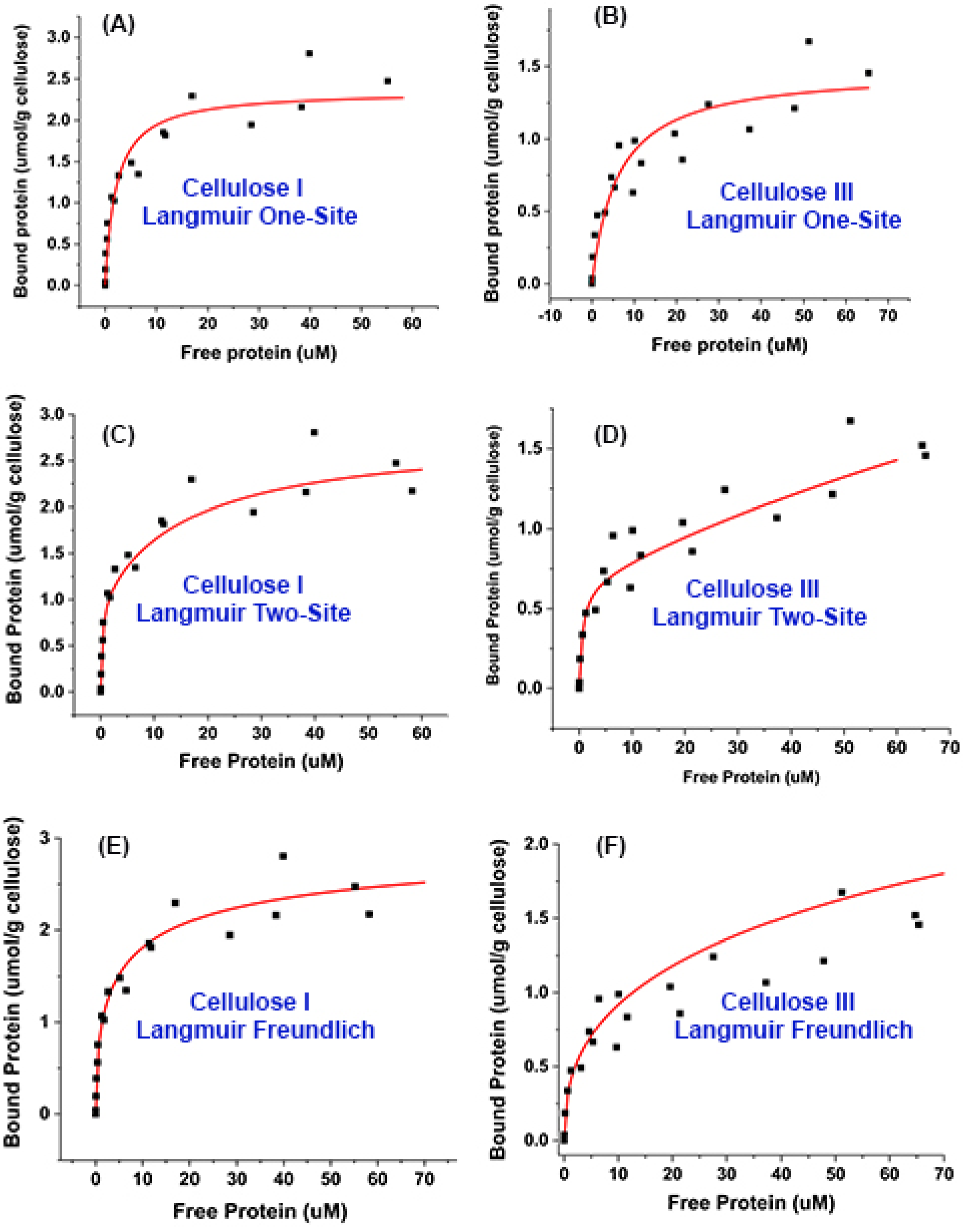

**Fig. S6.** AFM imaging of Cladophora-derived cellulose I or CI (A, D) and cellulose III or CIII (B, E) elementary fibrils. The black line in panel A and B corresponds to the height signal (forward and backward) shown in panel D and E. As is can be seen in panel D and E, there is no difference in the forward and backward signals, hence only the forward signal was used to analyze the height profile dimensions which are depicted in panel C. Cellulose I fiber showed a peak with a clear shoulder while no such shoulder was seen for the cellulose III fibril, which is consistent with the modification in fiber shape after pretreatment.

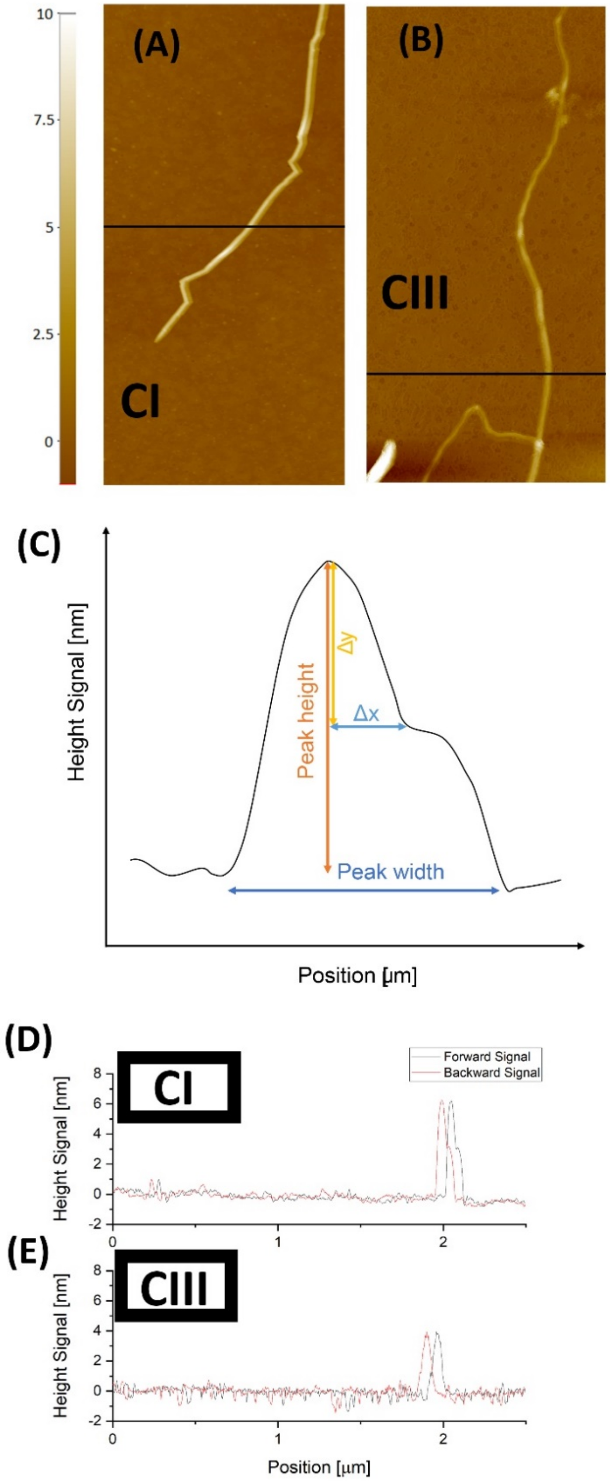

**Fig. S7.** Protein adsorption data and fitted partition coefficient slopes for various Type-A CBMs to Cladophora-derived Cellulose I (in black) and Cellulose III (in red).

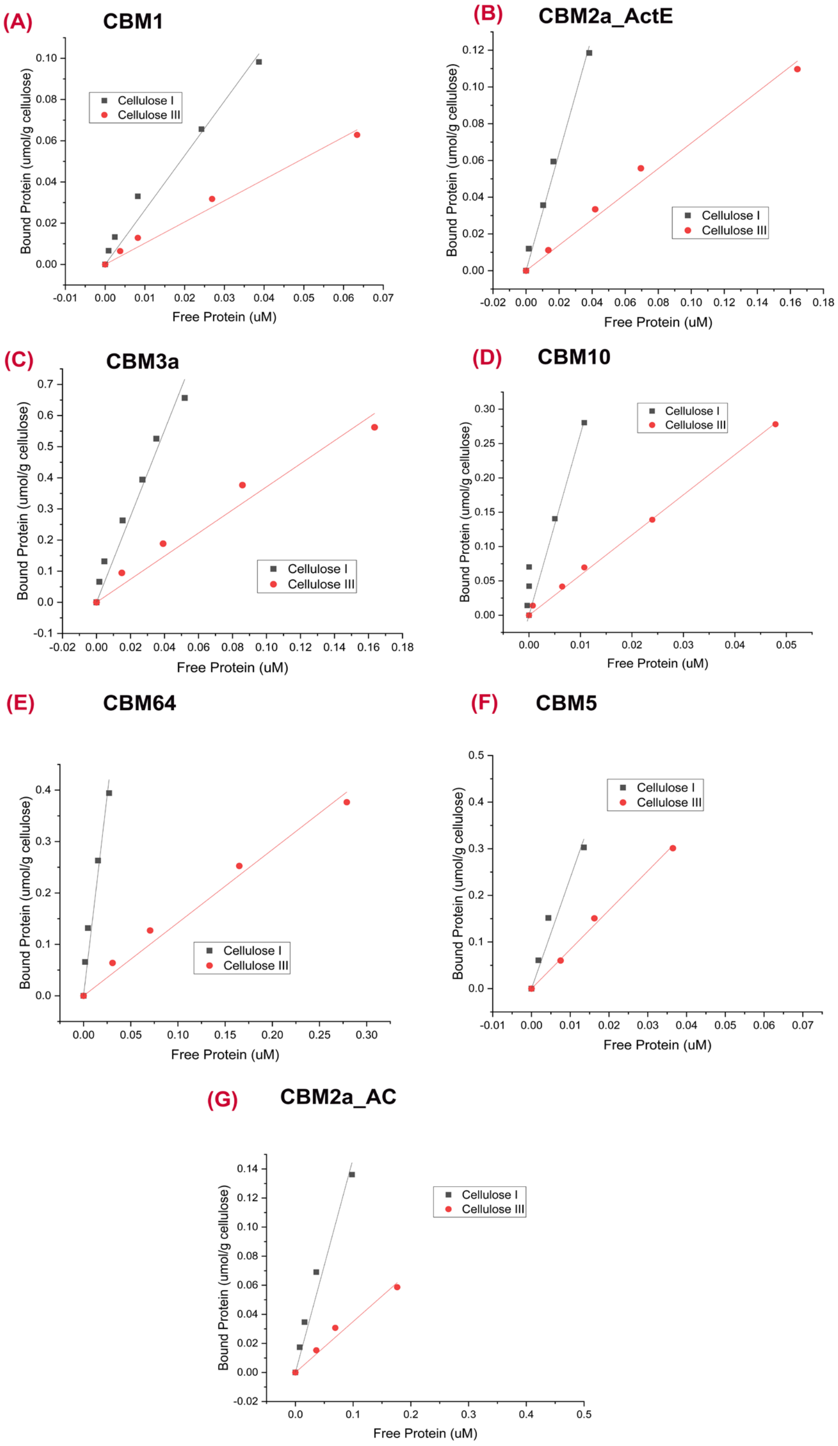

**Fig. S8.** Quartz crystal microbalance (QCM) based analysis of GFP-CBM3a binding parameters towards Avicel based cellulose I (CI) and cellulose III (CIII) nanofibrils. Representative QCM traces for binding/unbinding dynamics of GFP-CBM3a towards cellulose I (A) and cellulose III (B) are shown below. Representative binding model fits (in green) to raw QCM data (in black) is shown for binding (C) and unbinding (D) regimes. Model details are provided in the SI appendix methods section. Finally, relationship between energy dissipation parameter to QCM frequency is shown over the course of the GFP-CBM3a QCM binding assay (E).

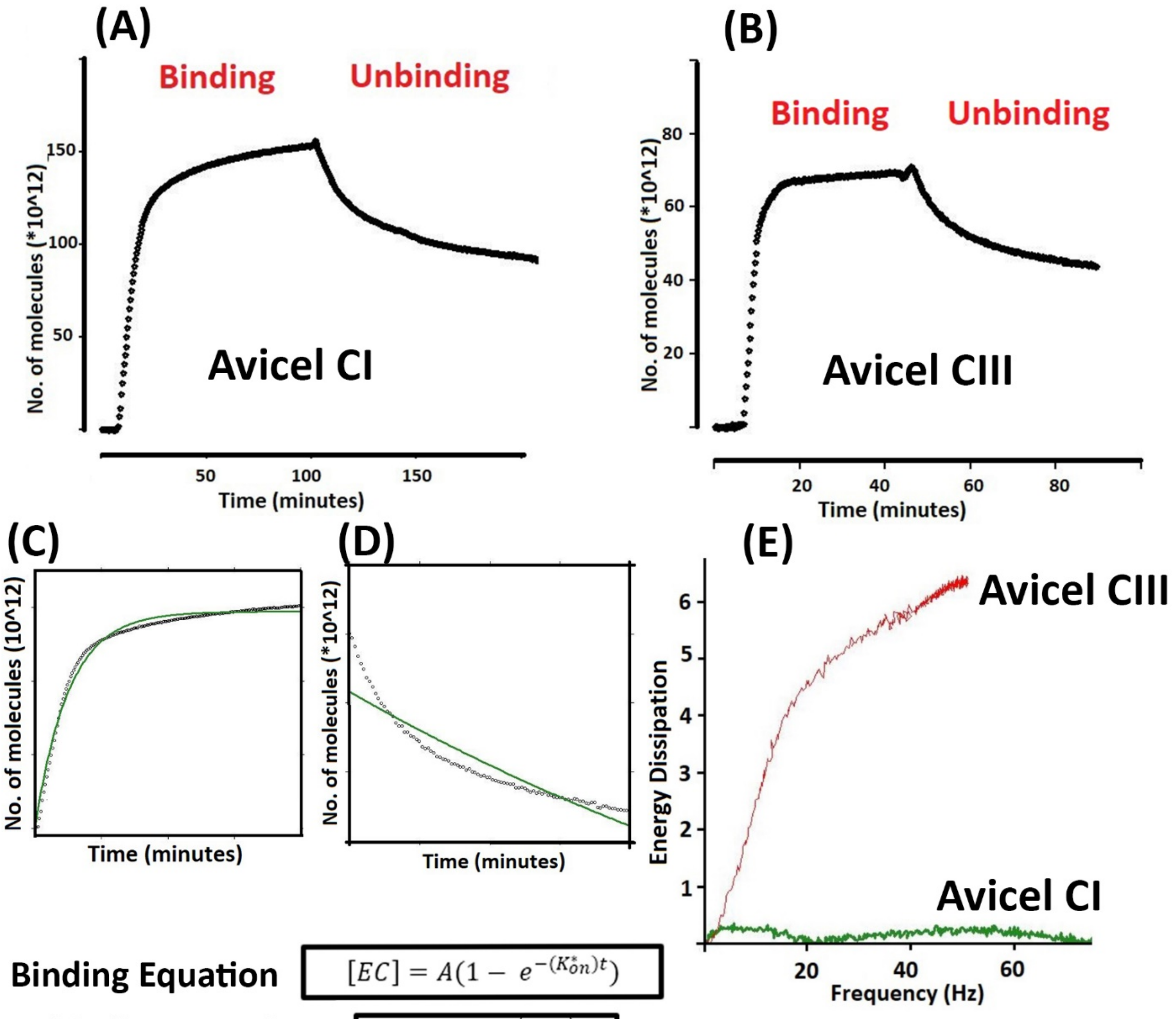


$$[EC] = A(1 - e^{-(K_{on}^{*})t})$$

$$[EC] = Ae^{-(K_{off})t}$$

**Fig. S9.** Characterization of GFP-CBM3a binding to Cladophora CI/CIII using FRAP. Panel A-C show snapshots of the FRAP image acquisition with panel A as the image before photobleaching, B first frame after local photobleaching and panel C at the end of the image acquisition. Panel D and E show example recovery curves for cellulose I and III respectively. Note that the shown curves are baselined to zero as mentioned in the SI methods section. Panel F and G compare the off-rate and $F_M$ histograms for GFP-CBM3a on cellulose I (CI) vs. cellulose III (CIII) with the Gaussian fit parameters (mean ± s.d.) as insets.

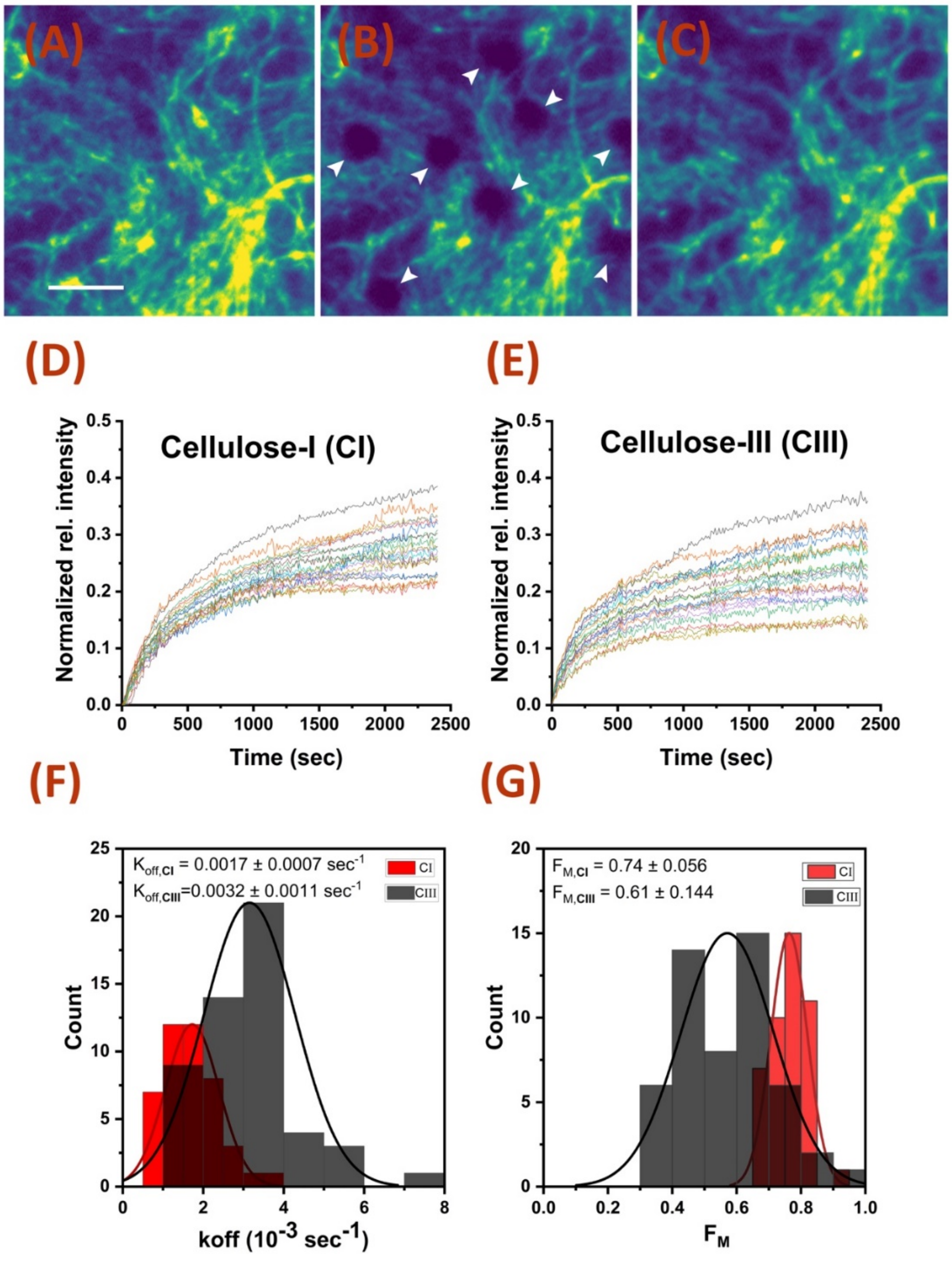

**Fig. S10.** (A) Force vs lifetime raw data scatterplot for the CBM1 bond on Cladophora celluloses I (blue) and III (brown). Total number of events measured (N) on cellulose I is 410 and on cellulose III is 214. For visual clarity, we omitted data points above 20 pN or 12 s from the scatterplot (31 for cellulose I; and 5 for cellulose III). We did not exclude any data from our report or analysis. Our one-way ANOVA test (B) concluded that there was no significant difference between the two entire datasets or at 2.5 pN intervals. Such a wide variance in lifetimes further supports our claim of multiple binding regimes.

**(A)**

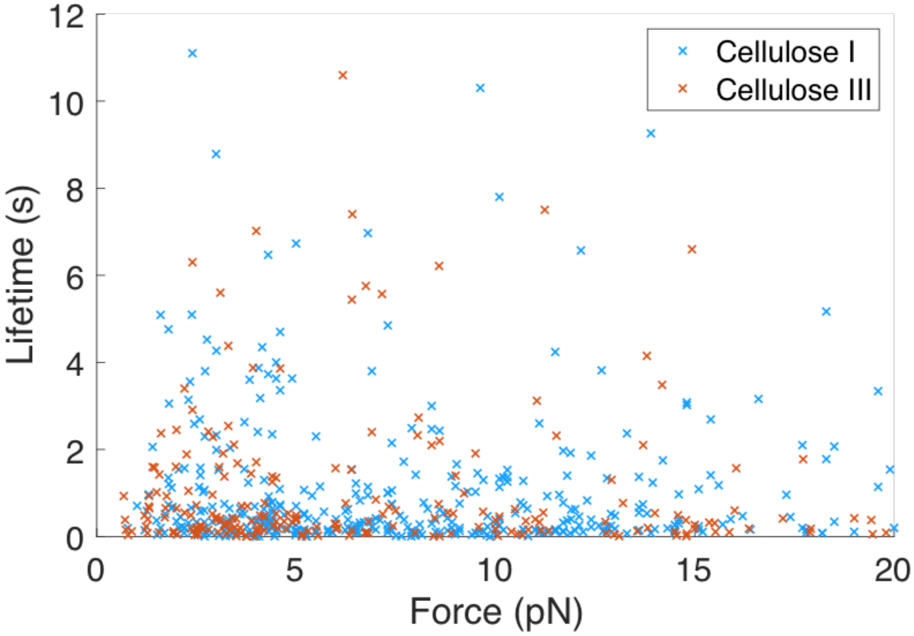


**(B)**

| Force Range | ANOVA Result |
|---|---|
| All Forces | $F(1,536) = 1.67$, $p = 0.20$ |
| 0-2.5 pN | $F(1,78) = 1.91$, $p = 0.17$ |
| 2.5-5 pN | $F(1,178) = 0.84$, $p = 0.36$ |
| 5-7.5 pN | $F(1,60) = 0.04$, $p = 0.85$ |
| 7.5-10 pN | $F(1,71) = 0.96$, $p = 0.33$ |
| 10-12.5 pN | $F(1,70) = 0.16$, $p = 0.69$ |
| 12.5-15 pN | $F(1,52) = 0.02$, $p = 0.93$ |
| 15-17.5 pN | $F(1,21) = 1.17$, $p =0.29$ |
| 17.5-20 pN | $F(1,20) = 1.82$, $p =0.19$ |

**Fig. S11.** Force vs lifetime relationships for the CBM1 bond on Cladophora derived cellulose I (blue) and filter paper (green) derived cellulose microfibrils. Both distributions failed to converge to the classical slip bond model and revealed that the CBM1-cellulose interaction is multimodal across different native cellulosic substrates. The reported mean lifetime of the CBM1-filter paper cellulose bond (3.03 ± 0.37 SEM) is higher than that of the CBM1-cladophora cellulose bond (1.41 ± 0.20 SEM).

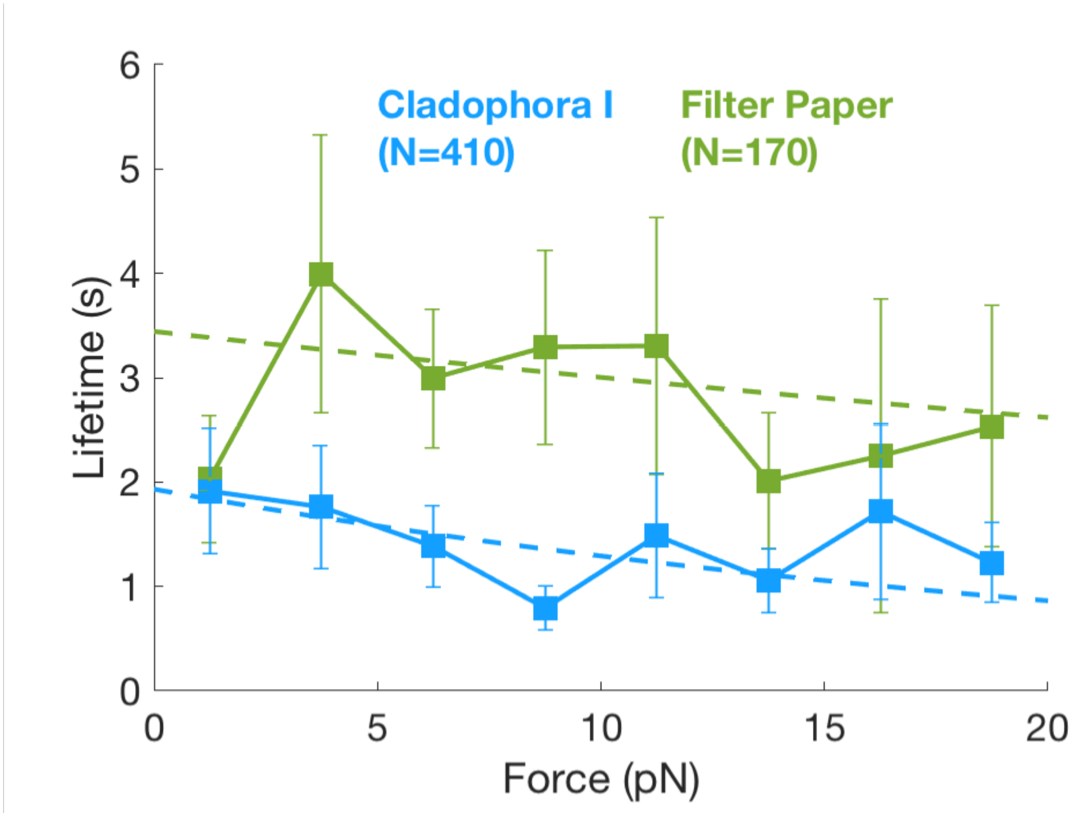

**Fig. S12**. (A) Force vs lifetime raw data scatterplot for the CBM1 bond on *Cladophora* cellulose I using an anti-His Fab in the assay construct (blue) and using full anti-His antibody (brown). Total number of events measured (N) using the full antibody is 233 and using the Fab is 187. For visual clarity, we omitted data points above 20 pN or 12 s from the scatterplot (8 for full antibody; 23 for Fab). We did not exclude any data from our report or analysis. Our one-way ANOVA test (B) concluded that there was no significant difference between the two entire datasets or at 5 pN intervals. Because of the statistical similarity, we combined both datasets to represent our CBM1-cellulose I data.

**(A)**

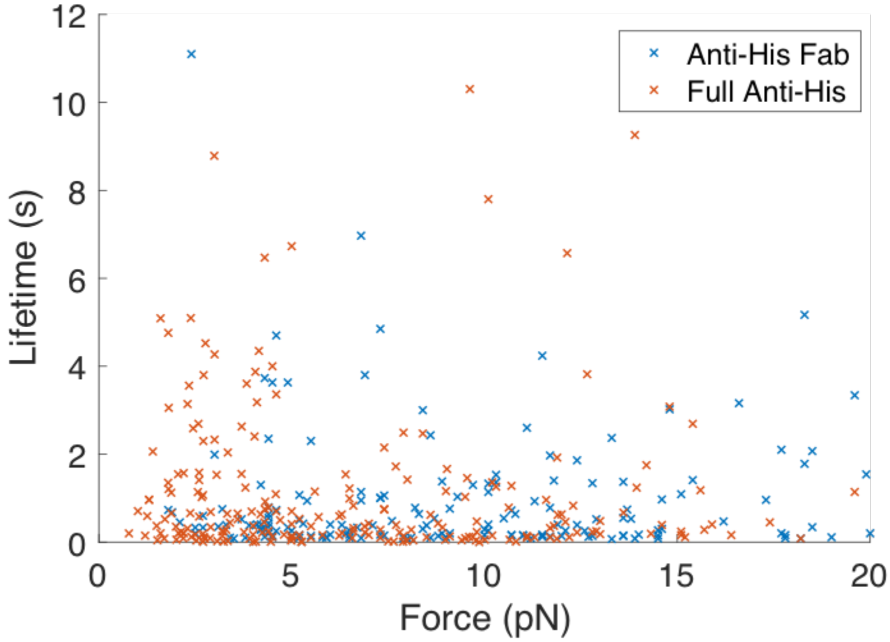


**(B)**

| Force Range | ANOVA Result |
|---|---|
| All Forces | F(1,370) = 2.74, p = 0.10 |
| 0-5 pN | F(1,149) = 0.13, p = 0.72 |
| 5-10 pN | F(1,104) = 1.10, p = 0.30 |
| 10-15 pN | F(1,86) = 3.00, p = 0.09 |
| 15-20 pN | F(1,25) = 0.13, p = 0.72 |

**Fig. S13.** (A) Force vs lifetime raw data scatterplot for the CBM1 bond on *Cladophora* cellulose I using the full CBM1 protein (blue) and using the Y31A CBM1 mutant (brown). Total number of events measured (N) using CBM1 is 410 and using the mutant is 93. For visual clarity, we omitted data points above 20 pN or 12 s from the scatterplot (31 for CBM1; 11 for Y31A-CBM1). We did not exclude any data from our report or analysis. Our one-way ANOVA test (B) concluded that there was no significant difference between the two entire datasets or at the 0-5 pN, 5-10 pN, and 15-20 pN ranges. However, there was a significant difference at the 10-15 pN range indicating the structural change does affect the CBM1-cellulose bond. Future single molecule studies could explore the effects of other protein structural changes on bonding.

**(A)**

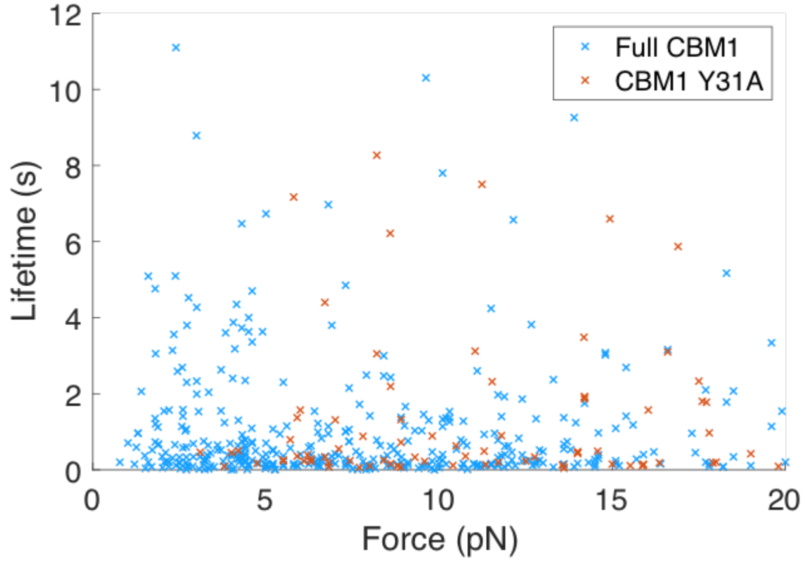


**(B)**

| Force Range | ANOVA Result |
|---|---|
| All Forces | $F(1,455) = 2.36$, $p = 0.13$ |
| 0-5 pN | $F(1,154) = 0.34$, $p=0.56$ |
| 5-10 pN | $F(1,139) = 2.44$, $p = 0.12$ |
| 10-15 pN | $F(1,115) = 5.66$, $p = 0.02$ |
| 15-20 pN | $F(1,41) = 0.13$, $p = 0.71$ |

**Fig. S14.** (A) Side-view and cross-section snapshot view from CBM1-cellulose unbiased MD simulations conducted for various distinct cellulose ultrastructures; cellulose I and cellulose III (with or without constraints that give rise to varying degrees of surface chain order or crystallinity index/CrI). See SI Movies S1-S3. Arrow direction indicates the preferred canonical binding orientation of CBM1 for cellulose I. (B) Average diffusivity of CBM1 on cellulose allomorph surfaces was estimated from the trajectories of the unbiased MD simulations to show significantly higher values for cellulose III versus cellulose I, again suggestive of weaker CBM1 binding interactions with the former substrate. (C) Steric clashes of planar CBM1 binding surface aromatic residues with cellulose III surface provides an atomistic basis for reduced binding towards cellulose-III. Root mean square fluctuations (RMSF) values seen for planar surface tyrosines (Y5, Y31, and Y32) was significantly higher for cellulose III of decreasing crystallinity.

(A)

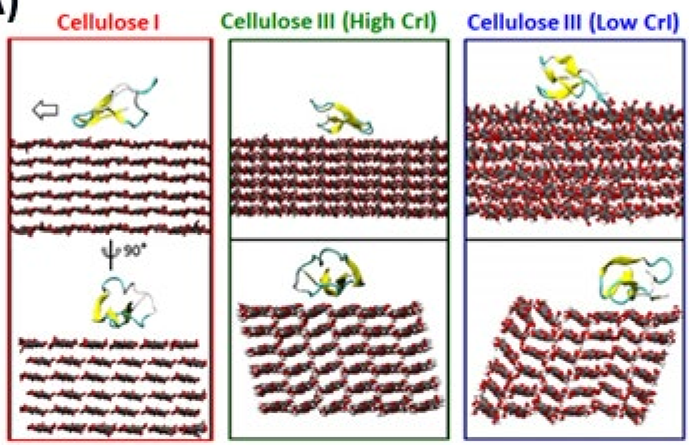


(B)

| CBM1 Diffusivity $(cm^2/s)$ | Cellulose I | Cellulose III (High CrI) | Cellulose III (High CrI) |
|---|---|---|---|
| ρ | $1.4 \times 10^{-11}$ | $6 \times 10^{-9}$ | $2.3 \times 10^{-5}$ |

(C) Root mean square fluctuation (RMSF) of key residues

| CBM1 Residue # | Cellulose I | Cellulose III (Low CrI) | Cellulose III (High CrI) |
|---|---|---|---|
| Y5 | 0.44 | 0.65 | 0.9 |
| Y31 | 0.47 | 0.54 | 0.9 |
| Y32 | 0.35 | 0.4 | 0.6 |

**Fig. S15**. Langmuir-type adsorption data (symbols) or model (dotted lines) fits (A) and Scatchard-plot representation of adsorption data (B) for Calcofluor White dye binding data to Cladophora-derived Cellulose I (in red) and Cellulose III (in grey). Here, the dotted lines represent Langmuir-two site model fits.

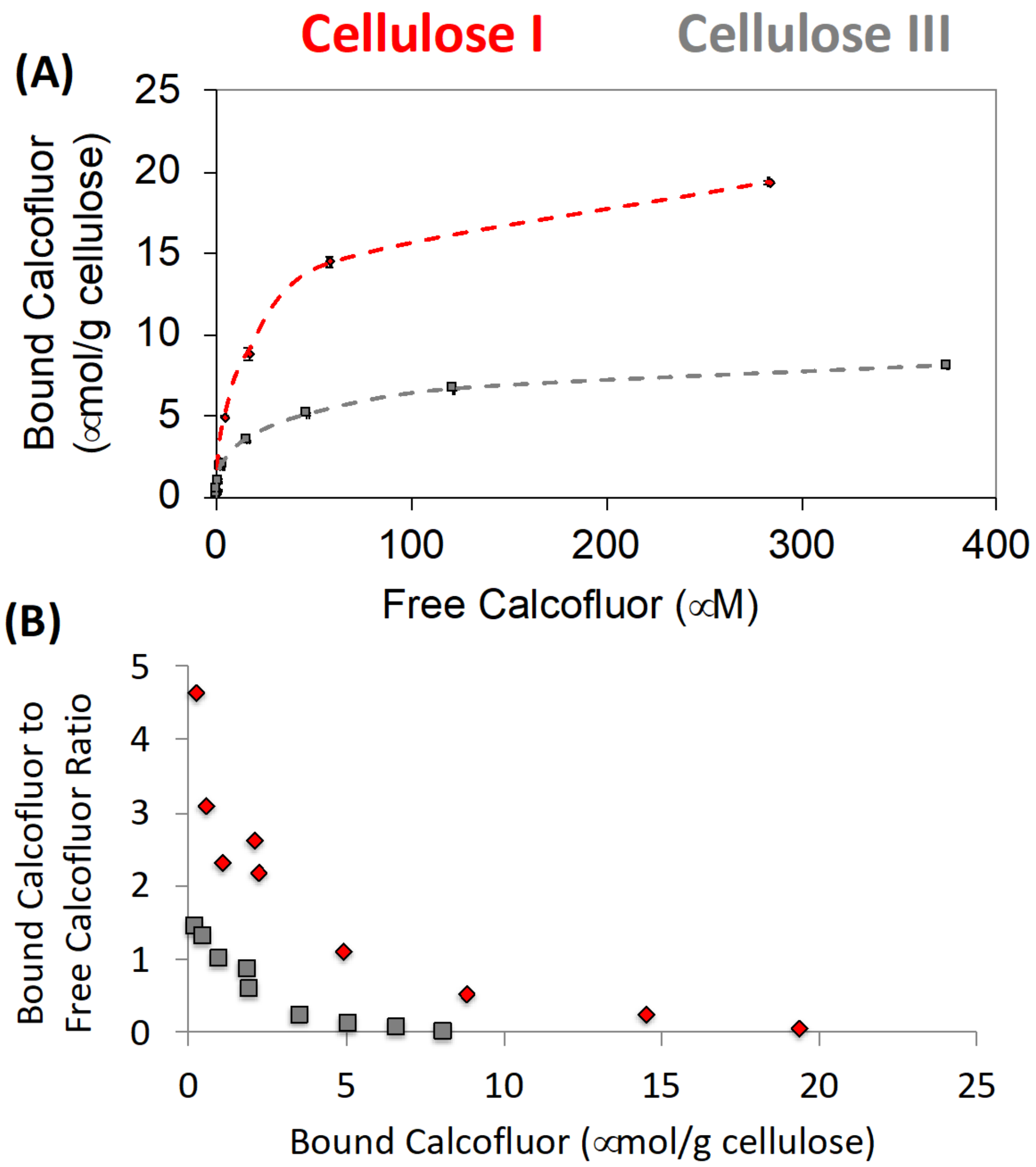

**Fig. S16.** Buffon needle model inspired CBM-cellulose binding orientation geometrical probability model overview/simulation methodology (A-E). Here, fractional events of wild-type (WT) and mutant (Y31A) CBM1 needle crossing along single ($\alpha$ ratio) or across multiple ($\beta$ ratio) cellulose chains is highlighted in (F), as predicted by the Buffon CBM/needle model.

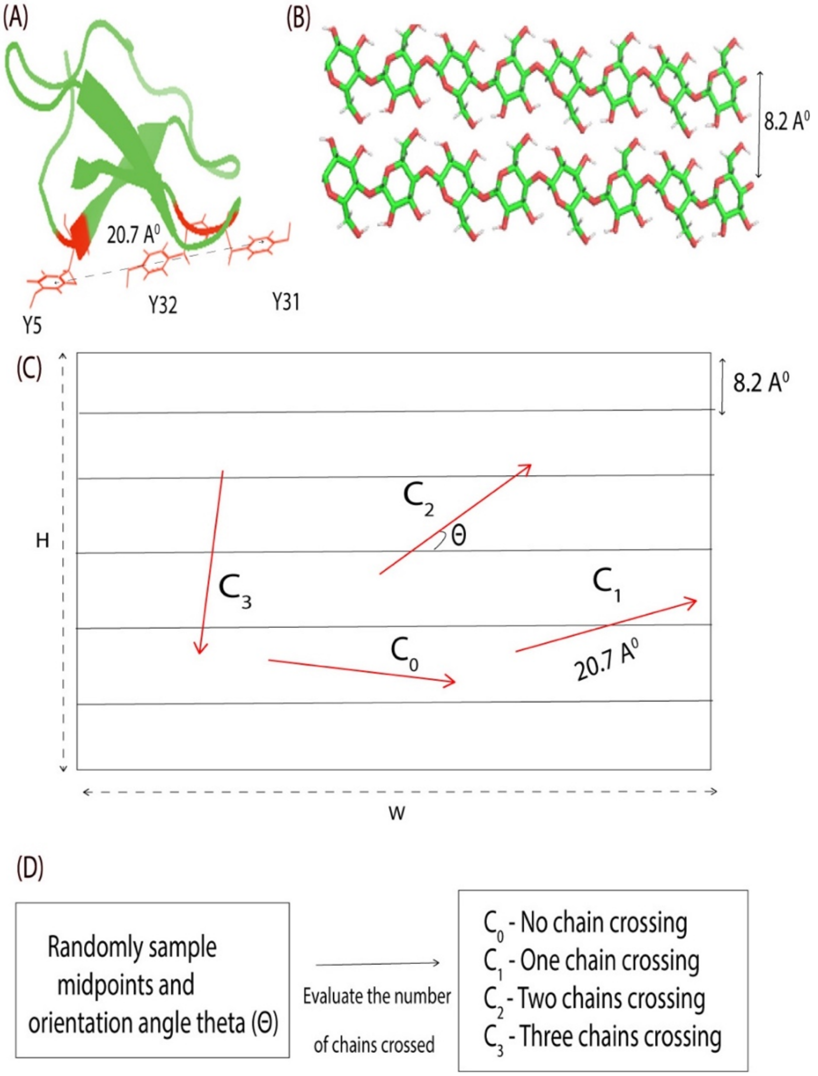


(E)

$$\alpha = \frac{C_0 + C_1}{C_0 + C_1 + C_2 + C_3}$$

$$\beta = \frac{C_2 + C_3}{C_0 + C_1 + C_2 + C_3}$$

(F)

| | α | β |
|---|---|---|
| CBM1 WT | 58% | 42% |
| CBM1 Y31A | 90% | 10% |

**Table S1.** Kinetic rate constants for GFP-CBM3a binding/unbinding towards nanocellulose fibrils estimated from QCM-D binding assay data are shown here. See SI appendix methods and results section for details.

| | A (molecules) | $K_{on}$* ($sec^{-1}$) | $10^{-3}$*$K_{off}$ ($sec^{-1}$) |
|---|---|---|---|
| Cellulose-I | 145.55 $\pm$ 0.40 | 0.13 $\pm$ 0.02 | 4.60 $\pm$ 0.21 |
| Cellulose-III | 97.71 $\pm$ 1.62 | 0.14 $\pm$ 0.01 | 11.30 $\pm$ 0.04 |

**Table S2**. Langmuir one-site model fitting results for truncated datasets generated from CBM1 binding to Cladophora cellulose I and III allomorphs. Here, model parameters for the original data (with no truncation) and data truncated to 15 or 50 μM free protein concentrations are reported.

| Langmuir One-Site Binding Model (Without truncation) | | |
|---|---|---|
| | Cellulose-I | Cellulose-III |
| $N_{max}$ | 4.34 ± 0.05 | 3.32 ± 0.09 |
| $K_d$ | 8.69 ± 0.31 | 10.55 ± 0.91 |
| RMSE | 0.17 | 0.17 |
| Langmuir One-Site Binding Model (truncated to 50 uM) | | |
| $N_{max}$ | 3.95 ± 0.05 | 3.23 ± 0.14 |
| $K_d$ | 7.05 ± 0.24 | 9.88 ± 1.07 |
| RMSE | 0.008 | 0.16 |
| Langmuir One-Site Binding Model (truncated to 15 uM) | | |
| $N_{max}$ | 3.18 ± 0.07 | 1.82 ± 0.05 |
| $K_d$ | 4.68 ± 0.21 | 2.91 ± 0.21 |
| RMSE | 0.008 | 0.05 |

**Table S3.** Sensitivity analysis was performed by changing the predicted Langmuir one-site model parameters by 10% or 20% and checking for goodness of fit (i.e., root-mean square error or RMSE) of the original CBM1 binding dataset.

| Sensitivity Analysis for One Site Model | | |
|---|---|---|
| | | RMSE |
| Original Data | | 0.17 |
| 10% | $0.90N_{max}$ | 0.23 |
| | $1.10N_{max}$ | 0.17 |
| | $0.90K_d$ | 0.17 |
| | $1.10K_d$ | 0.17 |
| 20% | $0.80N_{max}$ | 0.35 |
| | $1.20N_{max}$ | 0.35 |
| | $0.80K_d$ | 0.2 |
| | $1.20K_d$ | 0.19 |

**Movie S1.** Snapshot of side-view (top) and cross-sectional (bottom) view of CBM1-cellulose I unbiased MD simulations.

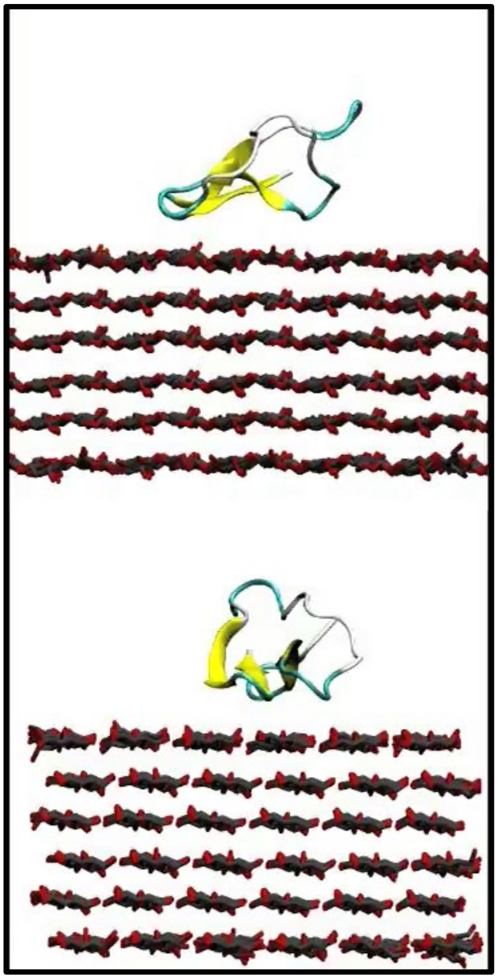

**Movie S2.** Snapshot of side-view (top) and cross-sectional (bottom) view of CBM1-cellulose III (high crystallinity) unbiased MD simulations.

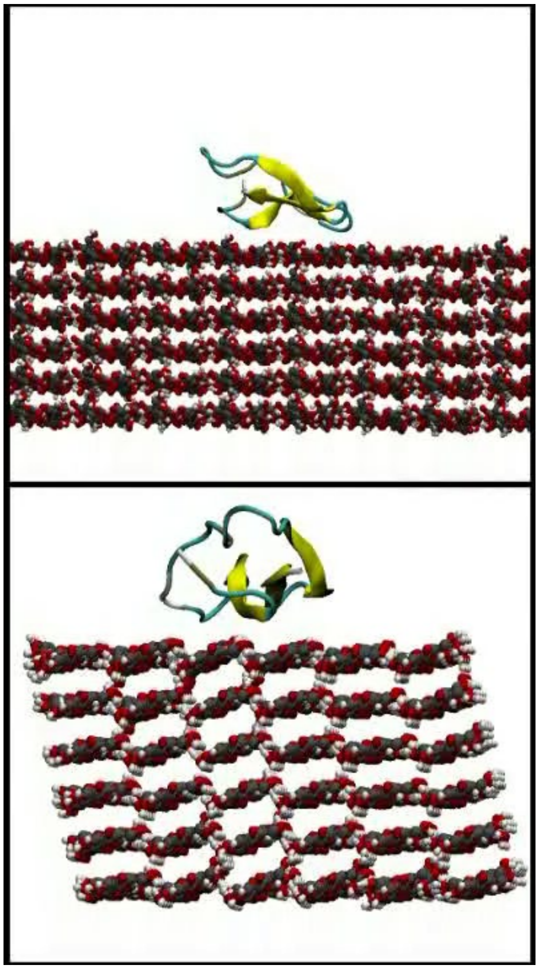

**Movie S3.** Snapshot of side-view (top) and cross-sectional (bottom) view of CBM1-cellulose **III** (low crystallinity) unbiased MD simulations.

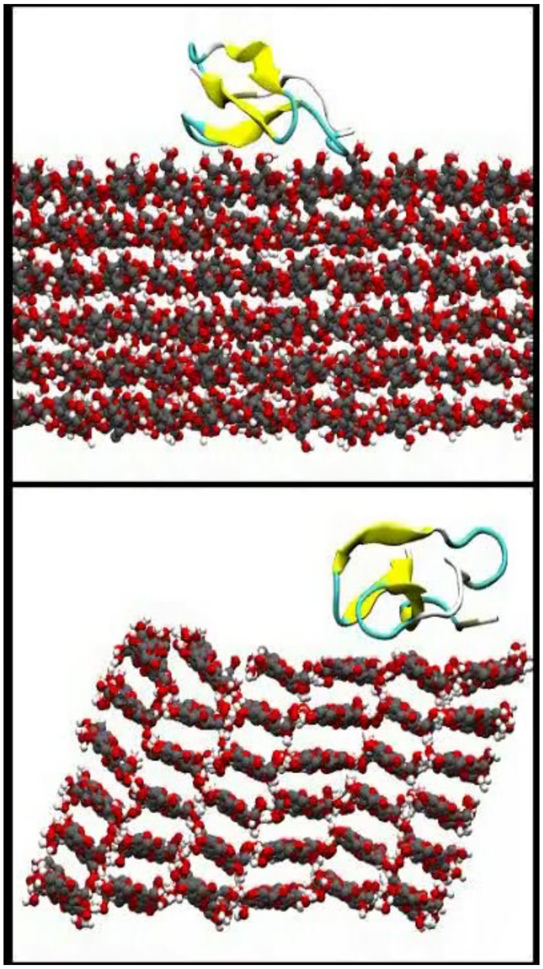
